# Micro-plaque assays: A high-throughput method to detect, isolate, and characterize bacteriophages

**DOI:** 10.1101/2024.06.20.599855

**Authors:** Gayatri Nair, Alejandra Chavez-Carbajal, Rachelle Di Tullio, Shawn French, Dhanyasri Maddiboina, Hanjeong Harvey, Sara Dizzell, Eric D. Brown, Zeinab Hosseini-Doust, Michael G. Surette, Lori L. Burrows, Alexander P. Hynes

**Affiliations:** Department of Medicine, McMaster University; Department of Biochemistry and Biomedical Sciences, McMaster University; Department of Chemical Engineering, McMaster University; Department of Pathology and Molecular Medicine, McMaster University; Farncombe Family Digestive Health Research Institute, McMaster University; Michael DeGroote Institute for Infectious Disease Research, McMaster University; School of Biomedical Engineering, McMaster University

## Abstract

The gold standard for the isolation and characterization of bacteriophages (phages), the plaque assay, has remained almost unchanged for over 100 years. The need for improvements to its scalability has been driven home by successes with personalized phage therapy requiring large phage libraries and rapid sensitivity testing. Using a robotic pinning platform, we miniaturized plaque assays from bacterial lawns to micro-colonies from 100 nl of inoculant, increasing throughput by >1000 fold without compromising sensitivity. A comparable manual workflow with one quarter the throughput maintained the same sensitivity. These micro-plaque assays can replace plaque assays as a new gold standard in phage biology. As proof of principle, we used our technique to isolate and de-replicate 21 unique *Pseudomonas aeruginosa* phages from a single environmental sample. We then demonstrated – using the same assay - that of 17 multi-drug resistant clinical *P. aeruginosa* strains, 15 were susceptible to infection by one or more of the 21 phages tested. Our method allows rapid isolation and de-replication of phages, as well as enabling screening of large phage libraries against bacterial isolates of interest.

## Introduction

Bacteriophages (phages) are bacterial viruses that, prior to the advent of genomics, were identified solely through their ability to lyse bacterial cells. This results in a drop in turbidity of a bacterial culture, or, more commonly, a zone of clearing in a bacterial lawn known as a ‘plaque’. Phages were first discovered using this type of plaque assay (Twort, 1915; D’Herelle, 1917), and it has remained the gold standard for isolation, quantification, and characterization of phages ever since (Clokie and Kropinski, 2009). Uniquely, plaque assays allow one to reproducibly count phages, distinguish between phages and other bactericidal entities, identify mixed populations of phages and enable their purification, and quantify the bactericidal efficacy of a phage against its host.

However, the plaque assay is laborious, beginning with a dilution series of phage lysate. It requires at least one Petri dish per dilution of lysate per host bacterium; more commonly, three plates per dilution results in the use of up to 18 plates to obtain one reproducible titre. This inconvenience explains the popularity of the lower resolution (Khan and Nilsson, 2015) “spot” test where small aliquots of phage dilutions are spotted on a bacterial lawn and the titre determined from the lowest dilution resulting in individual plaques (Clokie, M. R *et al.,* 2009). The loss of resolution is a trade-off for the scalability, as 4-6 phages can be titred on a single plate if all infect the same host. The constraints of the plaque assay and its variations have impacted fundamental research but are also increasingly problematic for therapeutic applications of phages. Tight phage-host associations – down to strain-level specificity (Hyman *et al.,* 2010; Moineau *et al.,* 2001; Kudva *et al.,* 1999) – limit the broad-scale application of any given phage.

A prominent phage therapy case study featured a patient treated with phages for an *Acinetobacter* infection (Schooley *et al.,* 2017). The impressive publicity that followed its success made it a catalyst for changes in public perception, backing, and regulatory approval. Treating the patient was an exercise in personalized medicine: culturing the causal bacterium, testing its sensitivity to a large library of phages hosted at the Naval Medical Research Center, tracking the emergence of phage resistance, and eventually using the emergent phage-resistant strain as bait to isolate new environmental phages to add to the cocktail (Schooley *et al.,* 2017). The case highlights the value of personalized medicine but also illustrates the challenges inherent in finding the “right” phage(s) in time to aid in clinical care.

As in most cases of personalized medicine, phage therapy faces a problem of scale; how do we efficiently build large libraries of phages that can rapidly be tested (and re-tested when resistance emerges) against unique bacterial isolates from each new patient? Both phage isolation and phage host-range profiling depend on plaque assays, and while this kind of personalized approach has been used to treat many different bacterial infections (Dedrick *et al*., 2019; Suh *et al.,* 2020; Gainey *et al*., 2020; Chan *et al.,* 2021; Duplessis *et al.,* 2019; Yang *et al*., 2020; Pirnay *et al.,* 2024), current techniques are ill-suited to the task.

Companies working to advance phage therapy acknowledge this limitation of scalability. One alternative adopted by Adaptive Phage Therapeutics (MD, USA) is called ‘host range quick tests’ (HRQT) (Adaptive Phage Therapeutics, 2019). The test involves combining a single bacterial strain from an infected patient with a large phage bank of pre-purified phages and monitoring bacterial growth in broth using a colorimetric assay. This assay is quick and scalable and has proven effective at matching phages to patients (Rao *et al*., 2022; Khatami *et al.,* 2021; Doub *et al*., 2020; Cano *et al*., 2021; Schooley *et al*., 2017). However, broth-based screens fail to quantify phage sensitivity (Abedon, 2016; Hyman, 2019). Furthermore, they cannot readily be used to build new phage libraries, as they cannot distinguish mixed phage populations, characterize phages from their plaque morphology, or enable the isolation and purification of single phages. These shortcomings mean that the plaque assay remains the gold standard in the field.

Attempts to increase the throughput of phage isolation aim to identify a wide diversity of phages as quickly as possible. A recent paper presented a high-throughput screen for rapid phage characterization using a robotic liquid handler. This method scaled the standard spot test to a 96- density assay with the allowance of doing ten at a time. Although the method was comparable to the gold standard, the resolution and specificity of testing was compromised (Dufour *et al.,* 2024). Olsen *et al*. developed a high-throughput screen where 94 samples are processed through an in- broth enrichment followed by filtration and spotting on a bacterial lawn (Olsen *et al*., 2020). This approach was successful in screening >500 environmental samples and isolating ∼309 phages for four bacterial host species (Olsen *et al.,* 2020). However, the approach has limitations. The ‘spot test’ (Mazzocco *et al*., 2009) approach where undiluted lysates are placed upon a bacterial lawn is limited to 48 lysates per bacterial host and cannot distinguish between phages and other bacteriostatic/bactericidal compounds such as bacteriocins produced by the enrichment host, nor can it identify ‘unique’ phages. Downstream direct plaque sequencing suggested as many as 38% (*E. coli*) of zones of clearing did not result from sequenceable phages, and as many as 66% (*S. enterica*) of the phages sequenced were not unique (Kot *et al.,* 2020; Olsen *et al.,* 2020).

Here, we describe a scalable and efficient approach to isolating and characterizing multiple phages for the preparation of therapeutic libraries. Miniaturizing the standard plate-based assay from a bacterial lawn to hundreds of individual colonies arising from dense 100 nl aliquots of host bacteria enables us to preserve some of the best features of plaque assays while increasing throughput by >3 orders of magnitude over the traditional format. We demonstrate how this method works in comparison to the gold standard across five model phages, and then exploit it in a custom workflow designed to rapidly generate and characterize large, diverse phage libraries for testing.

## Results

### Core Method Summary: Micro-Plaque Assay

We overlay 100 nl standardized bacterial cultures and phage lysates, exploiting automation for high throughput phage screening and detection as illustrated in Figure 1. The end product of each assay is an agar plate containing one or more bacterial hosts pinned at a 1536-colony density with a lysate overlay. Time-course imaging of the plate enables the collection of serial datapoints to construct growth curves – although for most applications, end-point imaging suffices.

**Figure 1.**
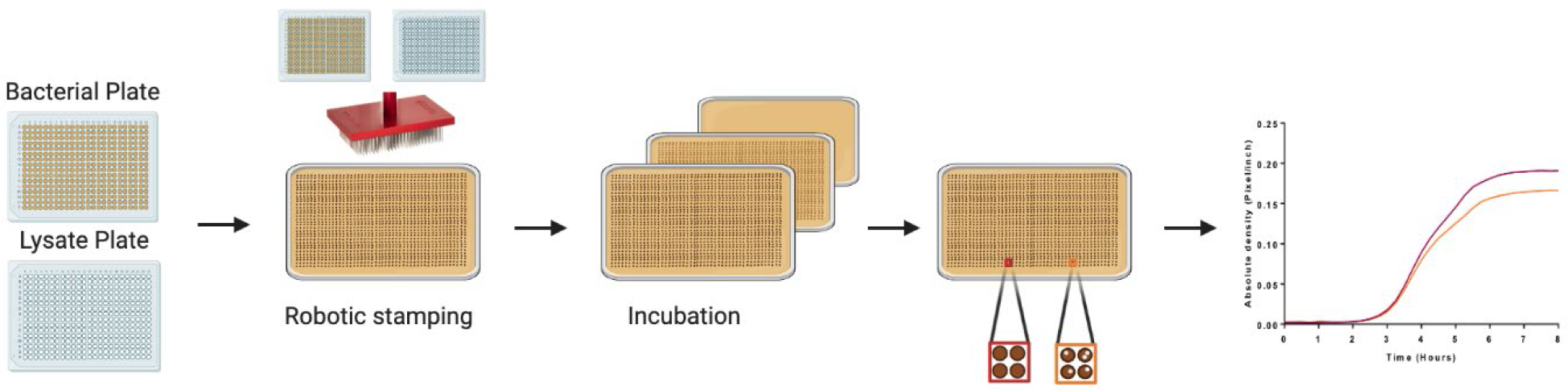
**Micro-plaque assays, an approach for high throughput phage screening**. Schematic demonstrates robotic bacterial stamping and overlay of phage lysates using the Singer Rotor HDA. Overlays are done at a 1536-per plate density. Plates were incubated for 16-18 h. Time-lapse images generated using a scanner every 30 min can be used to generate growth curves based on the absolute density of each colony formed.

### Validation of Micro-Plaque Assays

To validate our method, we used five well-characterized *Escherichia coli* phages spanning five different families; T3 *Autographiviridae* (Krüger & Schroeder 1981), T4 *Straboviridae* (Epstein *et al.,* 1963), HK97 subfamily *Hendrixvirinae* (Juhala *et al.,* 2000), M13 *Inovirida*e (Smeal *et al*., 2017 & Moon *et al*., 2015) and MS2 *Fiersviridae* (Davis *et al*., 1964 & Kuzmanovic *et al*., 2003) representing obligately lytic (T3, T4, MS2), lysogeny-capable (HK97) and chronic life cycles (M13). For consistency, all phage lysates were generated using the F’ *E. coli* K12 ER2738 (Table S1). These phages allowed us to explore the sensitivity and resolution of the micro-plaque assay relative to gold standard plaque assays.

We first carried out a traditional double agar overlay plaque assay on two different hosts for each phage (Figure S1), to obtain titres and morphology of plaques. These were then compared to micro- plaque assays of those same lysates arranged at 1536-density using a high-throughput screening Singer Rotor HDA pinning robot. Despite the 2 mm size of the resulting colonies, we observed individual plaques with morphologies characteristic of each phage (Figure 2). For example, phage T3 generates large plaques (Figure 2B), and this was recapitulated in the micro-plaque assays – as were the characteristically turbid plaques of M13. The results in Figure 2 are presented with the inclusion of black food colouring in the agar for additional contrast, which did not change the sensitivity significantly (*p-value* >0.281, paired T-test, Table S3). However, darker colouring made it easier to visualize the plaques and observe them earlier in the incubation period; 4 h instead of 6 h.

**Figure 2.**
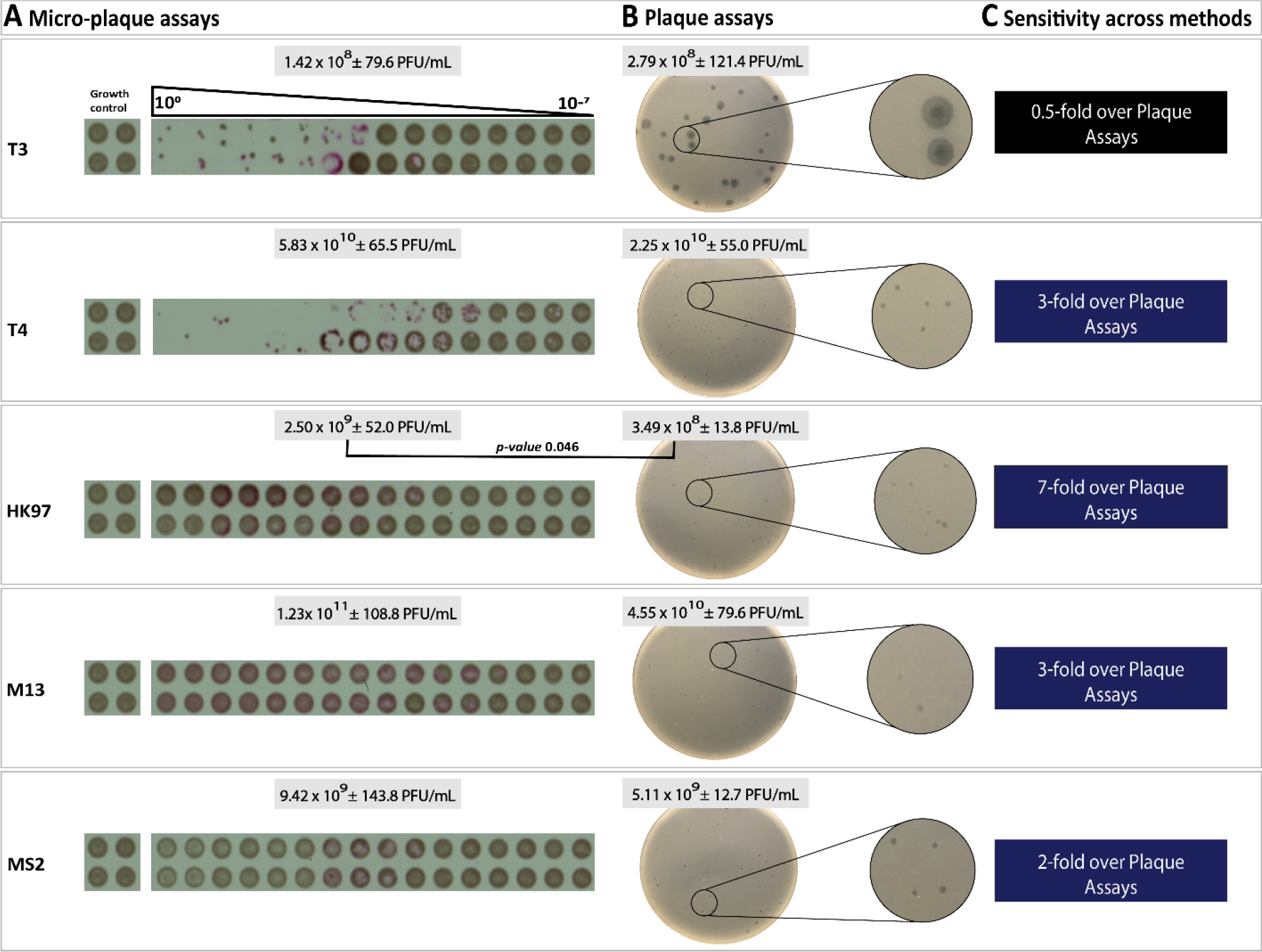
Plaque morphology and titration in both micro-plaque assays and the ‘gold standard’ plaque assays. A) Image from micro-plaque assays taken at endpoint from one biological replicate and their growth control in tetrad for all five model phages on *E. coli* K12 ER2738. Each micro-colony is represented by a tetrad of four technical replicates in a 10-fold dilution (10^0^-10^-7^). B) Image taken from a representative plaque assay. Titres shows the average of three biological replicates each of either 4 (micro-plaque assay) or 3 (plaque assay) technical replicates, ± standard deviation. C) Sensitivity across methods represented by fold change over plaque assay (PA). The fold-change was calculated with PFU/ml on average from the micro-plaque assay divided by PFU/ml on average from the plaque assays. All images were cropped from different plates and biological experiments, but not otherwise manipulated.

As with standard plaque assays, we count individual plaques and titre the phages in the micro- plaque assay across three biological replicates (Figure 2, S1, S2 and S3). The resulting titres were compared to those obtained with traditional plaque assays (Figure 2). For most phages, there were no statistical differences between the micro-plaque and traditional plaque assay titres, with the sole exception of HK97 (*p-value* 0.046, paired T-test), where there was a 7-fold increase in titre in micro-plaque assays compared to plaque assay (Figure 2).

To ensure these findings could be expanded beyond a single bacterial host strain, we also examined sensitivity of a second permissive *E. coli* host with same panel of phages – either *E. coli* B for T3 and T4, or a different K12 derivative (HER1382) for the other three phages. For the majority of the phages, there were no statistical differences in efficiency of plaquing (EOP) between the micro- plaque and plaque assay titres (Table S2). The exception was HK97, where both methods reported a significant change in EOP across hosts; the micro-plaques titre decreased 10000-fold, but the plaque assay titre decreased only 400-fold. On average across all model phages, the micro-plaque assays resulted in an increase in sensitivity of 2-fold over the gold standard.

While micro-plaque assays can be counted at endpoint after overnight incubation, an advantage of using time-lapse imaging is the fact that the density of each colony can be plotted to capture growth curves over the incubation period. The curves shown in Figures S1 to S3 were useful to calculate the optimal time to perform the micro-plaque assays. With our test phage and strain set, between 2 and 6 h is sufficient to determine the titre for phages plaquing on *E. coli* K12 HER2738 and *E. coli* B, whereas on *E. coli* K12 HER1382, between 4 and 9 h was sufficient (Supplement figures S1 to S3). In both cases, considerable time is saved when compared to traditional overnight incubations.

Acknowledging that access to a Singer Rotor HDA is limiting, we sought to replicate the robotic approach with one that can be performed manually. To stabilize the pads used to transfer phage/cells and ensure accurate overlay, we 3D-printed a holder for the Singer Pads (Figure S4). We also decreased the pinning density by four-fold to 384 microcolonies per plate. The resulting microcolonies generated using the 3D-printed Singer pad stabilizer are comparable to those obtained with the robot, although the lower density leads to more pronounced edge effects on the outermost colonies (Figure S5). While the process requires practice, the sensitivity was indistinguishable from that derived using the Singer Rotor HDA (Table S3). As the only effective trade-off is decreased colony density, this is a viable alternative to plaque assays that can reduce costs and requires no specialized equipment.

### Phind assay enables detection of a large number of isolated phages from a single environmental sample

Having demonstrated that our technique is a clear improvement over the gold standard, we sought to leverage its strengths. One advantage over other high-throughput phage assays is its capacity to detect and isolate individual phage plaques from environmental samples, exploiting the high- throughput platform to generate a large phage library. This assay to *find* phages we call phind. The high-throughput layout of the assay (Figure 3A) enables a maximum of 1536 independent host- phage pairings to be simultaneously tested.

**Figure 3.**
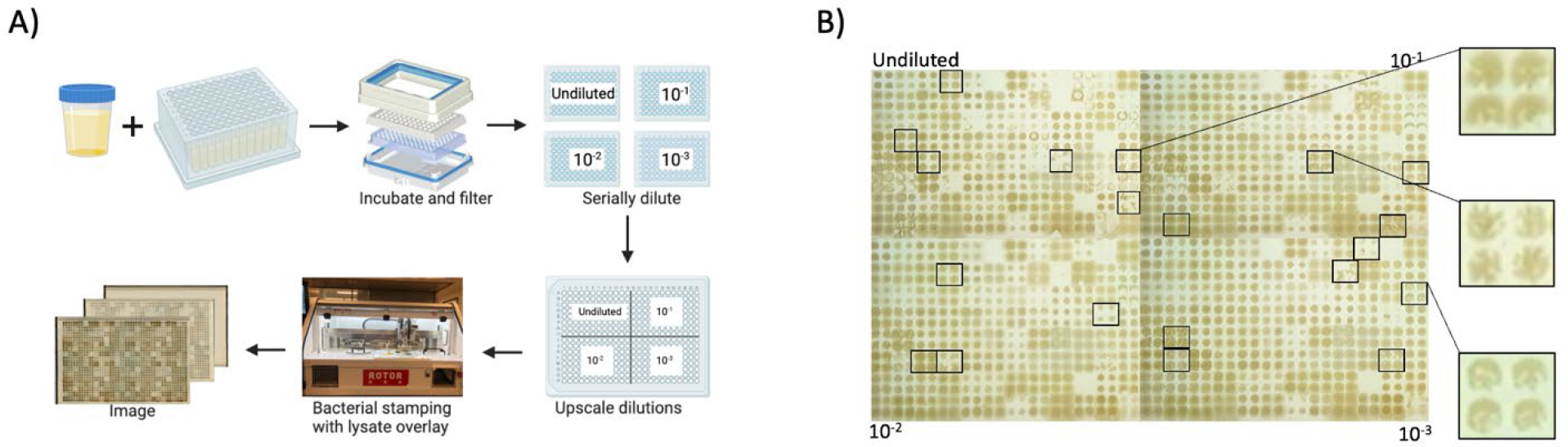
Phind assay designed to isolate phages from environmental samples. A) An environmental sample is enriched for phages by co-incubation with bacterial strains. The co-inoculated culture is incubated overnight and filtered with a 96 well vacuum manifold. The phage lysate plate is serially diluted, and each dilution plate is upscaled to 384 density, resulting in a single 384 plate containing 96 filtrates each at 4 different dilutions. A bacterial plate containing overnight cultures is prepared in a 96 well plate and then upscaled to 384 density, resulting in 4 replicates of each overnight culture. The Singer Rotor HDA uses 384 pin pads to stamp the bacterial plate first followed by an overlay from the lysate plate. The stamping is done 4 times within a single plate, totaling at a 1536-density containing 4 replicates per condition. Plates are incubated at 37° C. B) The Phind pipeline was used to identify new phages which were subsequently isolated. Areas of clearing following a phage-like dilution pattern, highlighted by the black boxes, were presumed to be phages and were picked for follow up.

To ensure reproducibility during the development phase, we limited our assays to 384 unique conditions, with each condition being tested 4 times. We used 96 different bait *Pseudomonas aeruginosa* host strains (Table S5.1) with enrichments on each plated at four different dilutions. The dilutions were necessary to differentiate between a distinct phage plaque and the effects of other growth-inhibiting bacterial products such as antimicrobial peptides, bacteriocins, or antibiotics (Fajardo, A, 2008; Björn, 2016; Soltani, S, 2021). With the resulting wealth of zones of growth inhibition, we focused on microcolonies where we could clearly observe distinct plaques. Plaques were identified as isolated zones of clearing where more concentrated samples yielded greater inhibition. As the primary goal of this assay was to obtain isolated plaques at endpoint, time-series imaging was not necessary. A total of 21 plaques (Figure 3B) were selected for further characterization to confirm that they were due to phages, and to establish the phages were unique.

### Phingerprint assay illustrates uniqueness of the isolated plaques

A major challenge when isolating large numbers of phages using high throughput methods – especially from a single environmental sample - is ensuring dereplication so the same phages are not repeatedly re-isolated. A similar issue occurs in the field of antibiotic discovery, where custom platforms have been developed for de-replication and adjuvant discovery (Cox *et al.,* 2017). We exploited tight phage-host associations and our high-throughput screen to generate a ‘host range’ fingerprint for each phage, allowing us to efficiently recognize if we have captured a phage similar to one previously isolated. We call this process phingerprinting (Figure 4A).

**Figure 4.**
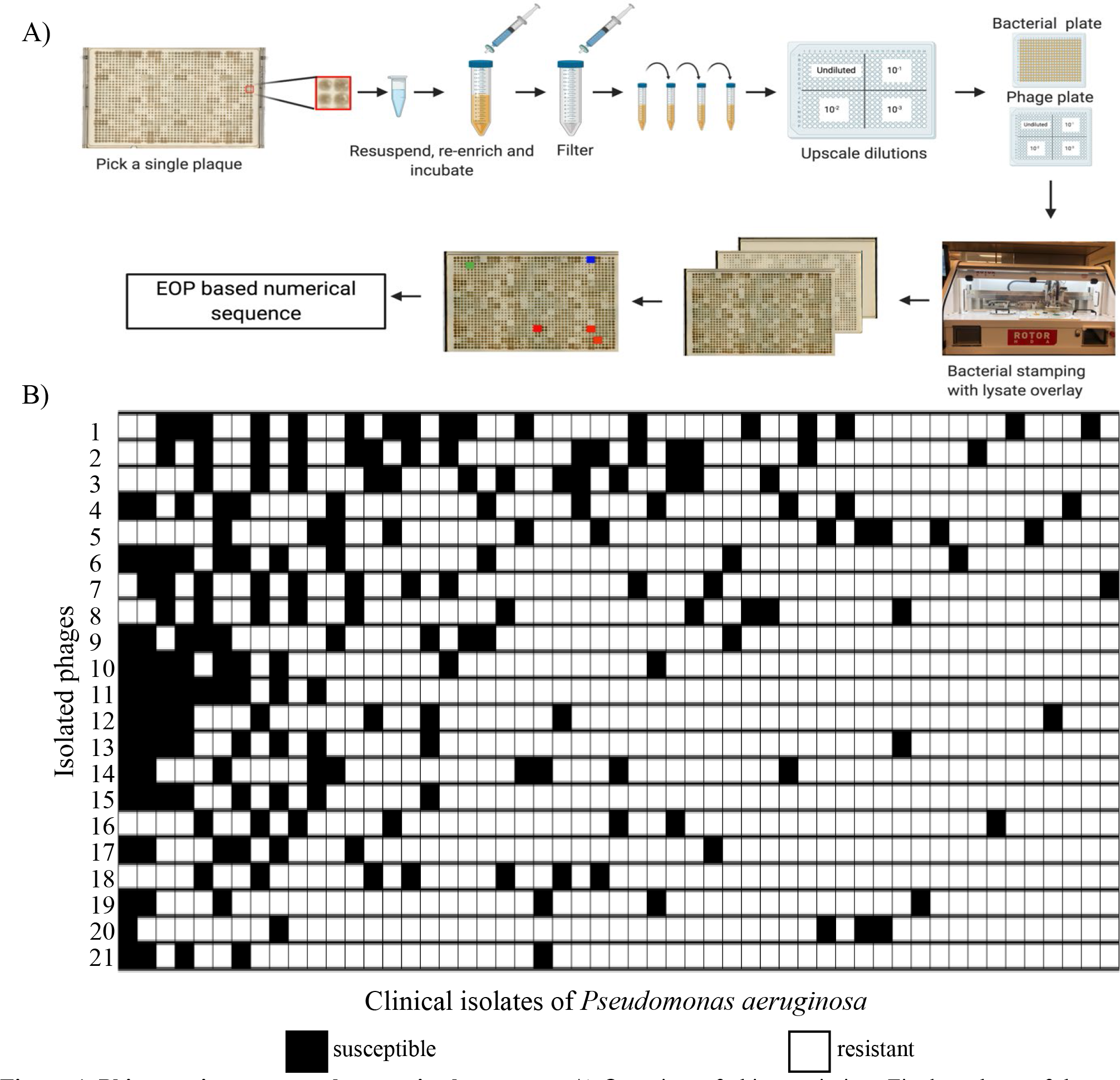
Phingerprint assay to characterize host range. A) Overview of phingerprinting: Final products of the phind assay are used to isolate single plaques. Plaques were picked and resuspended in phage buffer. The solution is then added to LB medium and inoculated with the original host. The co-inoculation is incubated at 37°C overnight, then filtered (0.45 μm PES) and serially diluted. The dilutions are upscaled to fit a 384 well format. This will result in a single 384 well plate containing one phage lysate at four different dilutions. A bacterial plate containing overnight cultures of 96 different strains is prepared. The Singer Rotor HDA uses 384-pin stamp pads to stamp the bacterial plate first, followed by an overlay of the lysate plate. The stamping is done four times within a single plate, totalling 1536 density containing four replicates per condition. B) A total of 21 different phages were generated via enrichments from a single environmental sample, each labelled with a unique phingerprint (host range). The uniqueness of each phingerprint is illustrated as a binary heatmap summarizing susceptible (lysis; black) or resistant (no lysis; white) across the 53/96 strains susceptible to at least one phage. Each column represents a different bacterial strain. Uninfected strains are not shown.

All 21 isolated plaques from our phind assay were characterized using the phingerprint assay, with the intent of sequencing only those that generated a unique phingerprint. The phingerprint was a characteristic host range pattern for that lysate on the 96 *P. aeruginosa* strains in our library (Table S5.2). As each was tested across four dilutions of lysate, the host range could also be reported as a numerical EOP for each host, rather than a simple presence/absence of lysis, should additional resolution be necessary. To minimize cost and labour, we opted to streamline the process by not further amplifying the isolated plaques to increase titre, further passaging on the host, or performing ultra-centrifugation and dialysis for increased purity. Accordingly, we assumed the phages were of low titre and thus a narrow range of dilutions (10^0^-10^-3^) was used to examine host range. Each of the 21 plaques resulted a different phingerprint considering presence/absence of lysis alone, meaning that their host ranges were unique (Figure 4B) even without considering EOP. To assess reproducibility, a single phage was also used to generate two phingerprint images under the same conditions. Both phingerprints were identical at the host range level, indicating high reproducibility (Figure S6).

Overall, we found that among the 21 isolated phages were those capable of infecting a total of 53 out of 96 bacterial strains – with the broadest infecting 17 strains, while the narrowest infected only five. Some hosts were susceptible to as many as 13 different phages, while others to only a single phage – and as such, served as valuable tools to de-replicate our phages. Phenotypically, all 21 phages generated a unique phingerprint. To ensure that the plaques were indeed due to phages and to determine their level of uniqueness, all were sequenced.

### Genetically Unique Phages

Each of the phages from the 21 unique phingerprints was sequenced, resulting in 45 metagenomic bins containing assembled sequences of >1500 bp and 5x coverage. Those bins were then sorted into phage “bins” which required a minimum of one hallmark phage annotation (capsid or terminase). From the 21 phage lysates we obtained 25 phage “bins”, even after subtracting all reads mapping to hosts to exclude any induced prophages. Due to use of non-purified lysates for phingerprinting, several preparations likely still contained multiple phages from the original environmental sample.

Rather than purify our phages from plaques, we instead simply amplified these low-titre samples in broth, enriching for whichever phage might out-compete the other in mixed samples. We obtained 19 samples yielding sufficient DNA to sequence, resulting in 21 phage “bins”, suggesting this single secondary amplification in broth was sufficient for our purposes (Fig 5A). This yielded a total of phage 46 bins (Fig S7) across both sequencing runs. In keeping with the notion of mixed samples, 8 were present (<95% sequence identity) only in the first, cruder preparation (Fig 5B). Across the two batches, 17 unique (<95% sequence identity) bins were shared, while 4 were unique. Of the 21 bins from our secondary amplification (batch B), 14 were considered unique (<95% sequence identity to any other), for a total of 22 unique (<95% identity to all others) bins from a single environmental sample.

**Figure 5.**
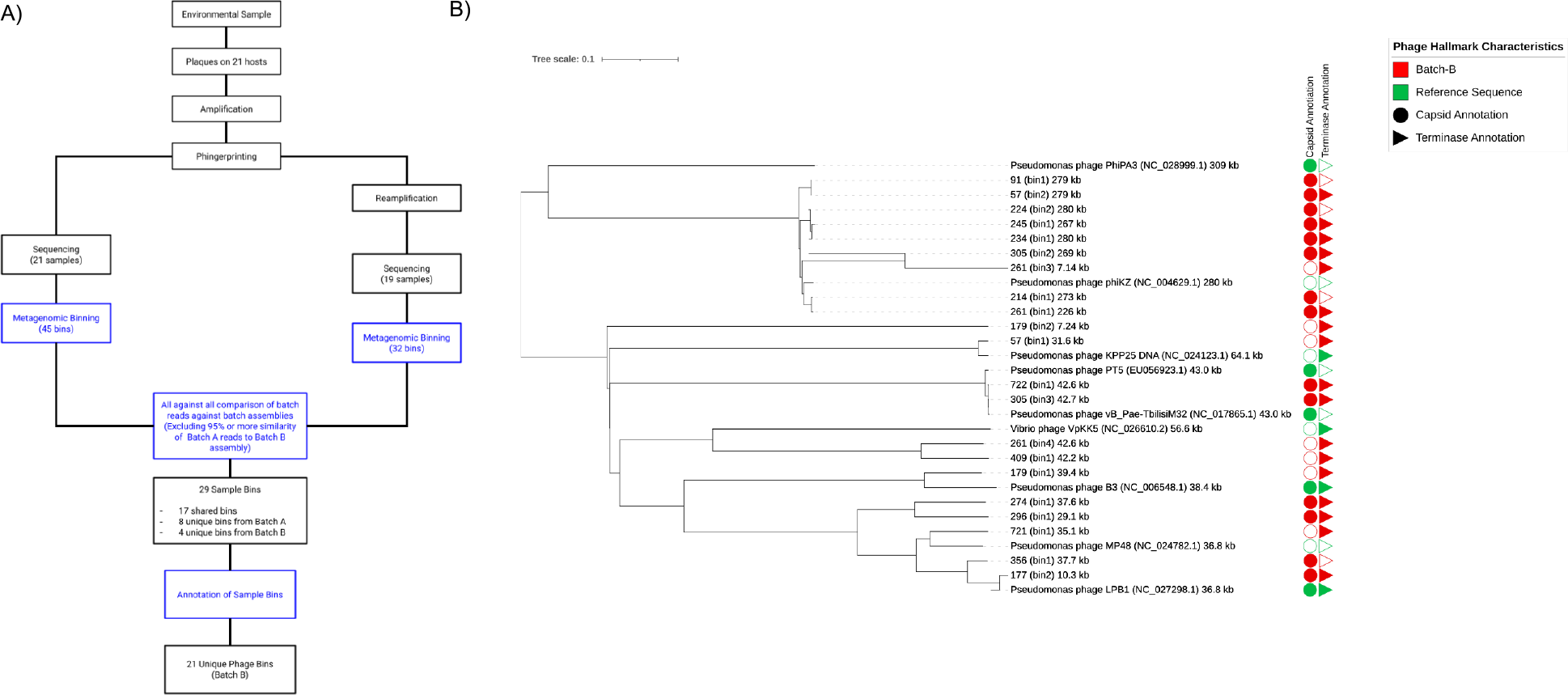
21 unique phages isolated from a single environmental sample. Identification of 21 phage bins using the Micro-plaque Assay workflow and an all-against-all genomic content comparison for batch A and batch B. A) A high-level overview of the Micro-plaque Assay process used to obtain the 21 unique phage bins. The process begins from environmental sample resulting in a phage bin containing genomic sequences of a sample as well as hallmark phage characteristics (capsid and/or terminase). The divergence in the process workflow between batch A and batch B was the addition of a purification step prior to the genomic sequencing for batch B. Identical phages were identified across batches by comparing genome length and annotation characteristics of phage. Unique samples across batches continued to downstream analysis. For any sample that was identical between the two batches, the sample from batch B was selected for downstream processing due to the additional purification step conducted. B) A proteomics phylogeny demonstrating the diverse number of phage bins identified by the Micro-plaque Assay created by VIPTreeGen Version 1.1.2. All samples from batch B (marked with an asterisk in panel B) are investigated. The samples compared are a minimum of 95% divergent from other samples within the two batches in experiment. Diversity of strains is shown by the red and green colours of the figure as the unique phage identified by the Micro- plaque Assay and the closest reference sequence available from the Virus-Host Database respectively. The two shapes associated with each sample (reference sequence or batch B sample) describe the presence or absence of two hallmark phage characteristics. The presence of a capsid (circle) or terminase (triangle) is defined by full or empty coloured shape.

We were intrigued to find several phages that were not considered genetically unique (>95% sequence identity) arising from plaques selected as unique in our dereplicating “phingerprint” assay. An example of this can be found in phages 305b_bin2 and 57b_bin2 (Fig S8). A trivial explanation of this is that the separate host range patterns are a result of the comparatively small (3%) sequence divergence between the two. However, it is also possible that it reflects host-controlled variation (Gencay *et al.,* 2019; Koskella *et al.,* 2013), and that the phage host range depends in part on the amplification host rather than being an inherent property of the phage. To test this, we chose phages whose host ranges overlapped, and amplified them on the shared host. One round of amplification was sufficient for host range convergence (Fig S9), confirming host-controlled variation, likely through DNA modification. This highlights that while phingerprinting can help identify different phages, identical phages may still appear unique based on the factors introduced by the amplification host– although this, too, is useful for understanding factors limiting host range.

To illustrate the diversity of phages selected by phingerprinting, we completed a whole-proteome phylogenetic tree using VIPTreeGen (Fig 5B) for only those 21 bins from our secondary, more pure amplification, including as references the closest related phage in the reference database (Virus-Host Database). With the exception of two distinct phages grouping most closely with a *Vibrio* phage (VpKK5), the most closely related reference phage was invariably a *Pseudomonas* phage, although these spanned everything from the transposable phages to the jumbo phiKZ-like phages. This analysis suggested that not only did we capture many different phages, infecting many different hosts, but we also captured phages across a very large phylogenetic space – all from a single environmental sample.

### Phage sensitivity profiling

Micro-plaque assays are effective for isolating many unique phages from even a single environmental sample, although it is not clear whether this would be advantageous for finding therapy-suitable phages. Of the 21 lysates generated from the phind pipeline, 18 were readily amplified and generated titreable plaques, and were therefore used to evaluate potential therapeutic application of the pipeline (Figure 6B). Each amplified phage lysate was tested on a set of 17 multi-drug resistant clinical isolates of *P. aeruginosa* to which they were naïve (i.e. were not a part of the phind or phingerprint libraries). In total, the 18 phages infected 15 of 17 isolates. The broadest host range phage infected 14 isolates, while the narrowest infected only one. Of the 18 phages, only one was incapable of infecting any of the clinical isolates tested. Some hosts were more susceptible to infection compared to others – with the most susceptible infected by 13 of 18 phages. These results indicate that this diverse set of phages isolated from a single environmental sample can infect multidrug resistant strains to which they are naïve. The phingerprint assay of these clinical isolates demonstrated distinct infection patterns, supporting the uniqueness of these phages.

**Figure 6.**
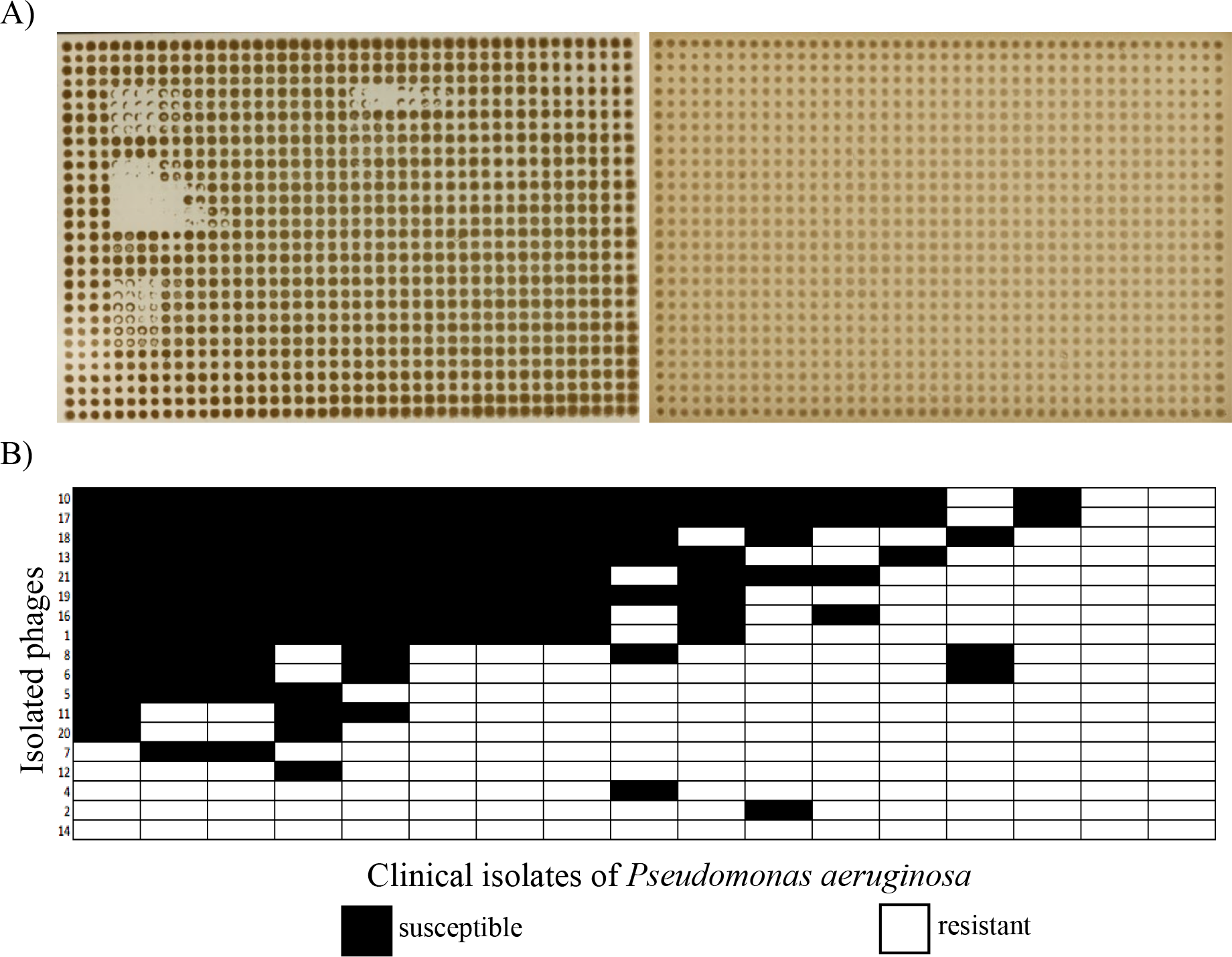
Phage efficacy on clinical *Pseudomonas aeruginosa* isolates. A) The left image illustrates plaquing of unique phages at different dilutions on a single clinical *P. aeruginosa* isolate. The right image illustrates an example where there was no visible plaquing by any of the phages on a clinical *P. aeruginosa* isolate. B) 17 clinical *P. aeruginosa* isolates were used to generate phingerprints for 18 phages. A phage’s ability to infect a host is illustrated as a binary heatmap summarizing susceptible (lysis; black) or resistant (no lysis; white). Each column represents a different bacterial strain.

## Discussion

Despite its century-long history of use for the isolation and characterization of phages, the plaque assay is too cumbersome for the high-throughput scalability demanded by modern applications such as phage therapy. Our alternative, a miniaturized version based on microcolonies generated using robotic overlays, shows comparable sensitivity across all phage families and hosts tested, massive increases in scalability even when phage and host are varied independently, and increased turnover. Each robotic overlay takes approximately a minute per plate, and results – with time-series imaging – can be obtained in as little as 6 h. The overall turnaround time of the assay outperforms that of traditional antibiotic susceptibility testing, which takes 16-24 h to complete (Goodwin *et al.,* 2007). We have also developed a parallel cheaper, manual version of the assay that has comparable sensitivity and 384-pin throughput, to ensure this micro-plaque assay can be used by labs across the world even in the absence of expensive robotics.

The scalability afforded by our assays enables quantitative phage sensitivity testing for clinical purposes, as well as the rapid isolation of unique phages to generate libraries to test against patient isolates. Some large phage libraries have been created, including the Naval Medical Research Center (Schooley, *et al*., 2017) with its 200 *Acinetobacter* phages, and the SEA-PHAGES (The Science Education Alliance Phage Hunters Advancing Genomics and Evolutionary Science) collection of over 10,000 *Mycobacterium* phages (Hatfull *et al.,* 2015), at times leveraged for treatment (Dedrick, *et al*., 2019; Hatfull *et al.,* 2019). However, establishment of these libraries has taken laborious efforts spanning many years. In contrast, with the ability to test up to 1536 independent hosts on a single plate, using environmental samples as a source of phages, we leveraged this assay to rapidly build phage libraries – in our proof of concept here, identifying 21 unique *Pseudomonas* phages from one sample on a single plate with our phind assay.

We can prioritize unique phages for further characterization by using our phingerprint assay to de- replicate phage hits. Although this assay does not guarantee uniqueness, as different phages could share the same host range, and similar phages could generate distinct phingerprints due to host-controlled variation, it can indicate which phages are more likely to be unique, allowing more efficient resource allocation. Scalability of operations could be further advanced using image analysis combined with machine learning to rapidly score images increasing the likelihood of identifying a unique phage (Faron, *et al*., 2016).

Finally, we show how we can use microcolony assays to screen clinical isolates for their sensitivity to a panel of phages, akin to an antibiogram. In our case, we showed that the environmental phages we isolated could lyse all but one of the clinical *Pseudomonas* isolates tested.

This entire process; phind, phingerprint and phage sensitivity profiling is enabled by our micro- plaque assay – a scalable, high throughput assay that preserves all the best features of the traditional plaque assay. If we solely relied on the traditional methods, the phind process described here would necessitate use of 96 petri dishes to detect phages across the bait library, and a further 96 for every 4-5 phages for host range testing (∼384), and another 17 for every 4-5 phages (∼68) for the antibiograms, not to mention the labour and time required. This would be unsustainable to incorporate into standard clinical care. We believe that the proposed HTS assay is a major advance over the century-old gold standard that, as its adoption by the field spreads, will enable prompt curation of libraries of diverse phages for any bacterial strain of interest advancing personalized medicine.

## Acknowledgements

We thank Félix R. Croteau for helping to generate the 3-D printed files for our Singer-free model and Rabia Fatima for helping to perform some of the phage titrations. We thank the Centre for Microbial Chemical Biology (McMaster University) for access to the Singer Rotor, Dr. Shirley- Anne Smyth and Stephen Teslic from the Waste Technology Centre at the Canadian Centre for Inland Waters for the environmental samples used for this work, as well as Dr. Gerry Wright for access to the strains from the Wright Clinical Collection at the Institute for Infectious Disease Research (McMaster University, Ontario) used for the phind and phingerprint assays. This work was funded by the David Braley Centre for Antibiotic Discovery, Farncombe Family Digestive Health Research Institute, and the New Frontiers in Research Foundation (APH: NFRFE-2018- 01441). LLB holds a T1 Canada Research Chair in Microbe-Surface Interactions (CRC 2021- 00311). ZHD holds a T2 Canada Research Chair in Bacteriophage Bioengineering, Ontario Early Researcher Award and a Discovery Grants Program.

## Author Contributions

APH, LLB, MGS and ZHD conceived of the study. DM and SF performed initial pilot work, with SF providing the imaging code and data analysis code, under the guidance of EDB. HH screened environmental samples. ACC performed validations and comparisons of the method against plaque assays, and the manual assay. GN performed the phind, phingerprint, sequence preparation, and final proof-of-principle on naïve strains. RSS performed all sequence analysis, with MGS contributing. ACC, GN, and APH wrote the manuscript, with all authors contributing editorially to the final product.

## Methods

### Model Phages & Bacterial Strains

The model phages and *Escherichia coli* strains serving as their hosts are listed in Table S1. Unless otherwise indicated, *E. coli* and *P. aeruginosa* strains were grown in LB broth at 37°C with shaking at 130 rpm, plated on LB 1% agar, 1.5% LB agar or added to molten 0.75% agar. To prepare stocks, each strain was inoculated into 10 mL of LB broth and incubated overnight at 37°C with shaking at 130 rpm. The following day, 1.5 mL of the overnight culture was centrifuged at 1300 *g* for 1 min. The supernatant was discarded, and the pellet re-suspended in 850 μL of LB broth and transferred into a sterile screw cap micro-tube with 150 μL of sterile glycerol and mixed. Bacterial strains were kept as a frozen stock at -80° C.

To obtain phage lysates a first amplification was made as follows. A scraping of bacterial strains (Table S1) from the frozen stock was inoculated into 10 mL of LB and incubated overnight. The next day, 100 μL of the overnight culture and a small scraping of each phage from frozen stock (Table S1) or from *P. aeruginosa* lysates obtained from the phingerprint pipeline (below), was added to 10 mL of LB broth and incubated until either lysis was observed or 18 h had passed – the latter was the case for all *P. aeruginosa* lysates. The inoculum was filtered with a 0.45 μm PES membrane. In order to increase titres, lysates of phages obtained from the phingerprint pipeline were subjected to a second amplification prior to being tested against other *P. aeruginosa* strains. Bacterial strains were inoculated into 10 mL of LB and grown to an OD600 of 0.2. One hundred μl of lysate was added and the culture incubated for five hours, then filtered (0.45 μm PES). All lysates were stored at 4° C.

### Environmental sample used for enrichment

The environmental sample used for enrichment was an influent wastewater sample collected on August 20^th^, 2019 from a site in Brantford, Canada graciously provided by the Water Technology Centre, Canada Centre for Inland Waters (CCIW), Burlington, ON. Bacterial plates for both assays were created using *P. aeruginosa* clinical strains from the Wright Clinical Collection (Table S5.1 and S5.2). The collection was taken as-is, with strains that had self-inhibited (filtrates generated from those strains that inhibited their own growth) replaced by strains highlighted in S5.2 to be used in the phingerprint assay.

*P. aeruginosa* grown in a plate for co-inoculation with environmental samples was grown in LB broth, at 37° C without shaking and scaled up to fit a 384-well format resulting in plates containing four replicates of each *P. aeruginosa* strain. Filtrates generated from these strains were then plated back on the host strain using the Singer. Strains which resulted in spontaneously induced prophages were removed in order to minimize background reading (Table S5.2).

### Plaque assays

Overnight LB broth cultures of the host bacterium were used to perform all experiments. Phages were first 10-fold serially diluted (10^-1^ to 10^-7^) in phage buffer (50 mM Tris-HCl pH 7.5, 100 mM NaCl and 8 mM MgSO4). To a tube of 3 mL of molten LB soft (0.75%) agar kept at 55° C, we added 300 μL of the bacterial culture and this was layered onto solid LB agar (1%). Once the soft agar was solidified, 5 μL of each phage serial dilution were spotted and left to dry for 10 min. Plates were typically incubated overnight at 37° C. The estimated titer from these spot tests was used to determine appropriate dilutions for full plaque assays. For these, 300 μL of culture and 100 μL of the appropriate phage dilution were added to 3 mL of molten LB soft agar and poured into a petri dish containing solid LB agar. The soft agar was dried then incubated overnight at 37 °C. Each dilution was tested in three technical replicates. After an overnight incubation, the dilutions which resulted in 30-300 plaques per plate were used to obtain the titer. The PFU/mL from plaque assays were represented by the average of the three biological replicates. A T-test was used to identify significant differences between the means of two groups.

### Singer Rotor HDA Preparations

Plus plates (Singer Instruments, cat. no. PLU-003) were prepared for Singer experiments following the protocol described in French *et al.,* (2020) with some modifications. Briefly, once the LB agar (1.5%) was at 55 °C, a small quantity (∼0.001 g) of black food colouring (Wilton, Item #610-981, US) was added aseptically and mixed well with the agar until it turned blue which increased contrast and allowed for easier detection of plaques. Addition of food colouring is not necessary if plaques are easily visible. Subsequently, 25 mL of the LB agar was transferred with a serological pipette into plates, careful to avoid bubbles. Plates were rocked against all edges to minimize surface tension. Plates were poured and left to dry on a carefully leveled surface. If not used immediately, plates were stored in bags at 4 °C and placed in the biosafety cabinet for 10-15 min prior to use to remove any condensation.

### Host Range determination/Efficiency of Plaquing

The efficiency of plaquing and host range determination were performed using the traditional double overlay plaque assay described above, and the Singer Rotor HDA assay. The Singer Rotor HDA was used to generate micro-plaque assays. A 384-well plate (Thermo Fisher, cat. no. 12565294) containing each strain (Table S1) was prepared under the same conditions as the plaque assays. An additional 384-well phage plate was prepared containing the five model phages, diluted as for the plaque assays. The rotor used 384 long-pin pads (Singer Instruments, cat. no. REP-003) to conduct a robotic overlay where the bacterial culture was stamped first, followed by the lysate. Each stamp was repeated four times, resulting in four technical replicates per condition and in a final 1536-colony per plate density. This was repeated for three biological replicates, each on its own plate. One pad was used to stamp all bacterial culture and pads were replaced prior to the stamp of each phage dilution. According to the manufacturer (https://singerinstruments.zendesk.com/hc/en-gb/articles/200756419-Use-of-compounds-with-the-ROTOR), Singer pads transfer on average 100 nl of liquid onto solid agar. This reference was used to calculate the titer from the micro-plaques. The PFU/mL from microcolonies was represented by the average of the three biological replicates. EOP was calculated using the formula: 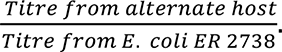. A paired t-test was used to identify significant differences between the means of two groups.

### Generation of Growth Curves

Flatbed color image scanners (EPSON Perfection V850 Pro, Model: B11B224201) were connected to a Raspberry Pi 3 (CanaKit, Model: B+) through a .SH file named *Code_scanner.sh* available in the following repository https://github.com/farncombephage/Micro-plaque_assays. Scanners were placed in a 37°C incubator or at 21°C and coded to automatically take images from Singer plates at given times within the duration of the experiment. Singer plate lids were removed and the plates were placed upside down onto the flatbed scanners. Images were acquired every 30 min over 16 h. All digital images were analyzed by FIJI (Schindelin *et al.,* 2012) to obtain absolute density of all micro-colonies using a macro called *Macro_time_series.ijm* and a file with the instructions to use the macro with time series images called *Instructions_macro_time_series_AC.txt* available in the GitHub micro-plaque assays repository. The absolute density obtained from all micro-plaques at all given point times were translated to growth curves using GraphPad Prism 7.0 (GraphPad software, Inc., CA, US). Final plots were created from each phage dilution and their no-phage growth control.

### Singer free-assay

We designed and printed an in-house Singer pad stabilizer to perform our micro-plaque assays at 384-density without using the Singer Rotor HDA. The file *3D_Singer_free_assay_file.gcode* is available in the GitHub micro-plaque assays repository mentioned above and was 3-D printed with the Ultimaker Cura S3 printer. The Singer pad was guided into the same position for stamping both the culture and the phage lysates. Once the holder was placed on top of the Singer plate, we use a double-sided tape to glue a plastic hanger on the top of the Singer pad to move the pad in and out from the Singer plate and perform both stamps. A general visualization of the 3-D printed Singer pad stabilizer is represented in Figure S4. An image was taken at the end-point of the micro- plaque assay at 384-density without using the Singer Rotor HDA is represented in Figure S5.

### Design of Phind and Phingerprint assays

#### The Phind assay designed to isolate libraries of phages

Fresh bacterial cultures were inoculated with the wastewater sample at a 1:10 ratio in a 96 deep well plate (VWR, cat. no. 10755-250). Any surface biofilms present following incubation were disrupted via agitation of the culture with plastic pipette tips. The plate was then centrifuged at 6000 x g for 10 min to pellet any debris that would hinder filtration. The supernatant – less 300- 500 μL to avoid the pelleted material – was transferred using a multi-channel pipette into a 96- well filter plate (Millipore Sigma, cat. no. MVHVN4525). A 96-vacuum manifold (Millipore Sigma, cat. no. MAVM0960R) was used to filter (0.45 μm PES) the cultures and resulting filtrates were collected in a 2 mL high-volume collection plate (Millipore Sigma, cat. no. MVCPN0025).

These filtrates were assayed as follows. Four sets of dilutions (10^0^ to 10^-3^) of the filtered lysates were created in standard 96 well plates (VWR, cat. no. 25381-056), then arrayed in quadrants of a single 384-well plate. The remaining lysates were stored at 4° C. A 384-well bacterial plate containing 96 overnight *P. aeruginosa* cultures and a 384-well plate containing 96 lysates at four different dilutions were used as the source plates for the assay. The Singer Rotor HDA used a 384 long pin pad to stamp the bacterial culture plate onto a prepared LB plus plate. The 384 well bacterial plate was stamped four times, creating a final 1536-density plate with four technical replicates per well organized in a tetrad. The 384 long pin pad was replaced with a new pad and the lysate plate was stamped on top of the bacterial inocula in the same manner. This set up resulted in conditions where each bacterial culture was paired with four dilutions of a filtrate generated from a co-inoculation of that same strain with wastewater, each in four technical replicates. The resulting plate was placed, lid open, in a scanner at 37°C and time-course data collected as per “Generation of Growth Curve”, above. This assay format was referred to as ‘Phind’.

#### The Phingerprint assay designed to characterize isolated phages and generate susceptibility profiles

The end product from the Phind pipeline was used to identify plaques. Plaques from the Phind plate were excised using a sterile 200 μl pipette tip, resuspended in 1 mL of phage buffer (50 mM Tris-HCl pH 7.5, 100 mM NaCl and 8 mM MgSO4), and incubated at room temperature for 15-20 min. Isolated phages were re-enriched by filtering (0.45 μm PES; Fisher Scientific, cat. no. 13100107) and adding the filtrate to 10 mL of LB broth inoculated with the host strain on which the plaque was originally isolated. Overnight cultures were centrifuged for 10 min at 6000 x *g,* with the duration of centrifugation occasionally extended depending on the quantity of biofilm formation. The supernatant was filter-sterilized (0.45 μm PES) and used immediately or stored at 4°C.

These re-enriched lysates were serially diluted 10^0^ to 10^-3^ and formatted to fit a 384-well plate with one lysate at four different dilutions. This was arranged so that each 96 well grid contained replicates of a single dilution. The above steps were repeated for each isolated plaque, creating a 384 well plate per isolated plaque containing four different dilutions. The 384-well bacterial plate containing 96 overnight *P. aeruginosa* cultures and the 384-well plate containing a single lysate at four different dilutions were used as the source plates for the assay. The Singer Rotor HDA used a single 384 long pin pad to stamp the bacterial plate onto all LB plus plates. After inocula were dried, a fresh 384 long pin pad was used per lysate plate to stamp on top of the bacterial cultures. All stamping was done four times, resulting in four technical replicates per condition. Plates were incubated at 37°C and time-lapse imaging was conducted, as above.

### DNA sequencing and annotation

Phage DNA was extracted from each lysate which resulted in a unique phingerprint using a standard phenol chloroform extraction with some minor additions (Sambrook & Russel *et al.,* 2001). Prior to the addition of phenol chloroform, 1 μl of DNAse I at 1 mg/ml (NEB, cat no. M0303L), 1 μl RNAse at 10mg/ml (NEB, cat no. T3018-1), and 1/10^th^ volume of DNAse I reaction buffer (NEB cat. no. B0303S) were added to 1 mL of phage lysate and incubated at 37°C for 30 min followed by a 60 min incubation at 65°C. Next, 10 μl of proteinase K at 20mg/ml (NEB cat no. P8107S) and a 2% final concentration of SDS was added and the mixture incubated for 1 h at 37 °C. The remaining process was as described in Sambrook & Russel (2001).

A NEBNext^®^ Ultra™ II DNA Library Prep Kit (New England Biolabs, cat. no. E7645S) protocol for Illumina was used to prepare DNA libraries for sequencing. It was optimized to be more cost effective with reduced volumes of reagents per reaction (Derakhshani *et al*., 2020). Briefly, the modifications made to the protocol provided by the NEB kit are as follows. In the DNA fragmentation step, the amount of genomic DNA, ultra-enzyme mix and ultra-buffer were reduced by 7.7, 6, and 4-fold respectively. For the adapter ligation step, 0.8 μL of adaptor (7 μmol) was added to the fragmented DNA. The amount of ligation master mix and enhancer added were reduced by 4-fold and 1.4 μL of USER enzyme and cutsmart buffer were added at a 1:1 ratio. ProNex® Size-Selective Purification System (Promega, cat no. NG2001) was used for bead purification/size selection. A 13.2 μL aliquot of beads was added to 12 μL of fragmented DNA. When preparing the library and attaching barcodes, all reagents and DNA added were reduced by 2-fold. A dual size selection was done to remove large fragments (above ∼1000bp) by adding 18 μL of ProNext beads to 20 μL of PCR product. The supernatant was then removed and small fragments (below ∼300bp) were removed by adding 4 μL of ProNext beads to the product. After incubation, fragments attached to the beads were washed and eluted. Resulting libraries were normalized based on DNA concentrations measured at 260 nm by Qubit 4 fluorometer (ThermoFisher, cat no. 33238). Sequencing was performed on a MiSeq platform with 2x300 reads.

The quality of reads was determined using FastQC v0.11.8 (Andrews, 2010). Reads were trimmed and assembled using Trimmomatic v0.38 (Bolger *et al.,* 2014) and SPAdes v3.13.0 [metaSPAdes mode] (Meleshko, 2017) respectively, both using the default settings. Quality assessment of draft contigs sequences was performed using QUAST v5.0.2 (Gurevich *et al.,* 2013). Host removal was performed for all samples for those hosts where genome sequences were available from the Wright lab clinical collection of 127 *P. aeruginosa* genomes. Alignment of host sequences to sample draft contigs using bowtie2 version 2.3.4.1 at default settings was performed and all unmapped reads were extracted by samtools version 1.7 using htslib 1.7.2 by the fastq command with the -f5 flag (Li, 2009). The unmapped regions of the alignment were reassembled using SPAdes v3.13.0 [metaSPAdes mode]. Following host removal, samples were grouped into metagenomic bins by MetaBat2 version 2 (Kang, 2019) with a minimum bin size of 5000 bp and a minimum contig size of 1500 bp, representing the s and m flags, respectively. The contigs of each metagenomic bin were concatenated into a single contig to represent the sample bin. These concatenated bin contigs were annotated by RASTtk v1.3.0 (Brettin, 2019) using the RASTtk default pipeline with the – ncbi-taxonomy-id flag set to the bacterial host strain 287 and the --domain flag set as Virus for the rast-create-genome command.

### Phage Profiling Against a Clinical Isolate

Phages isolated from the Phind step were amplified as described under “Model Phages & Bacterial Strains”. Dilutions from 10^0^ to 10^-7^ were performed and arrayed to fit onto a single 384 well plate. Seventeen clinical *Pseudomonas* strains näive to the phages were randomly selected from the Wright Clinical Collection. Serially diluted phages were robotically stamped onto a lawn consisting of a single bacterial strain. Plates were incubated for 18 h at 37° C on a scanner to collect time-course data.

## Supplementary information

**Figure S1.**
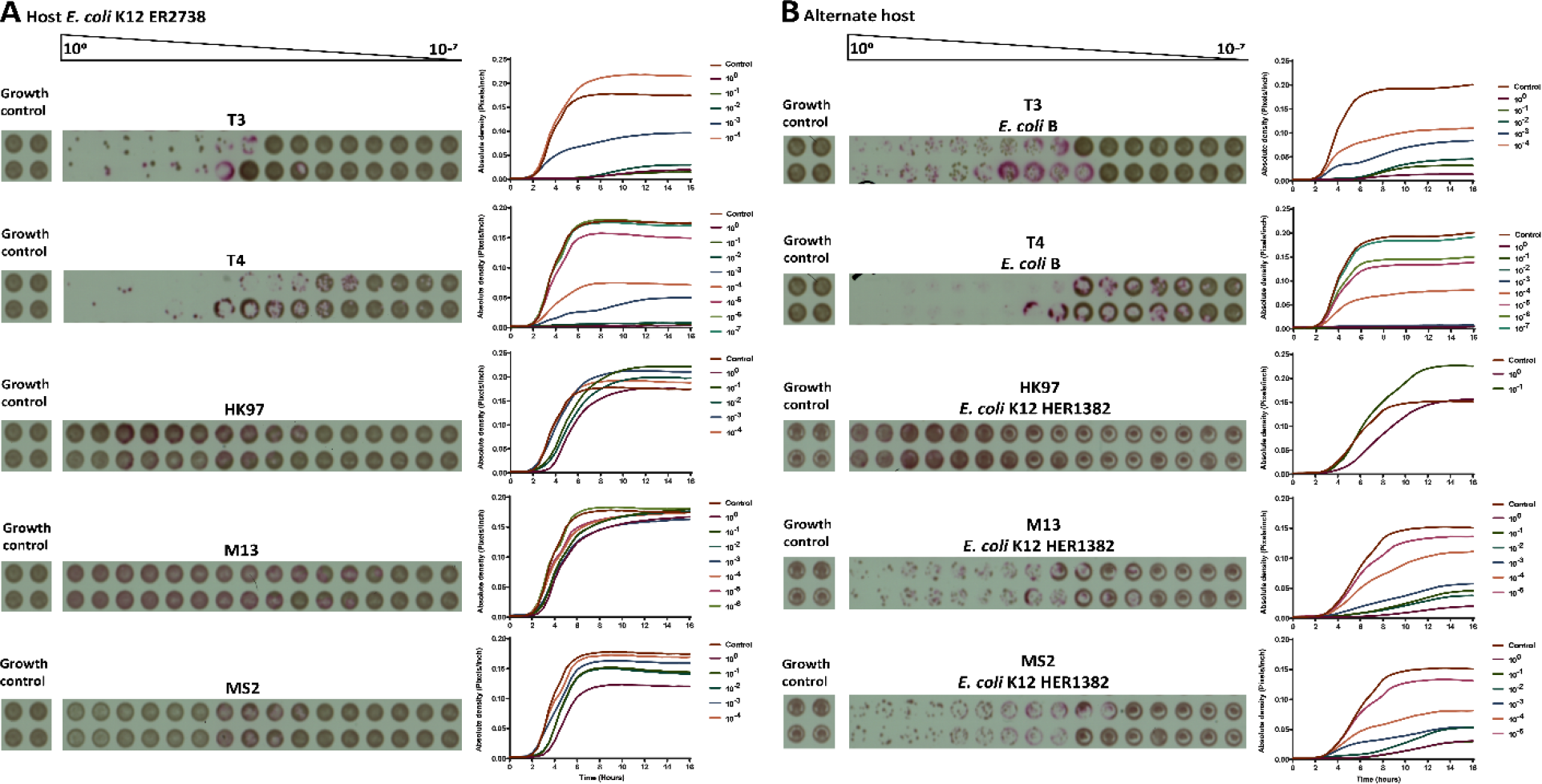
Biological replicate 1 of bacterial colonies with plaque formation of bacteriophages in micro-plaque assays. A) Left, image taken at endpoint for biological replicate 1. Each colony is represented by a tetrad of four technical replicates in a 10-fold dilution (10^0^-10^-7^). Growth controls for all phages was done using *E. coli* K12 HER2738 as host. Images were cropped from different plates according to the host, but not further manipulated. On the right side, plots represent the growth curves and their growth control, showing only curves in which plaques were counted, each curve represents the average of the four technical replicates per dilution. B) Same as A) but with different hosts as follows: T3 and T4 on *E. coli* B, with HK97 M13 and MS2 on *E. coli* K12 HER1382.

**Figure S2.**
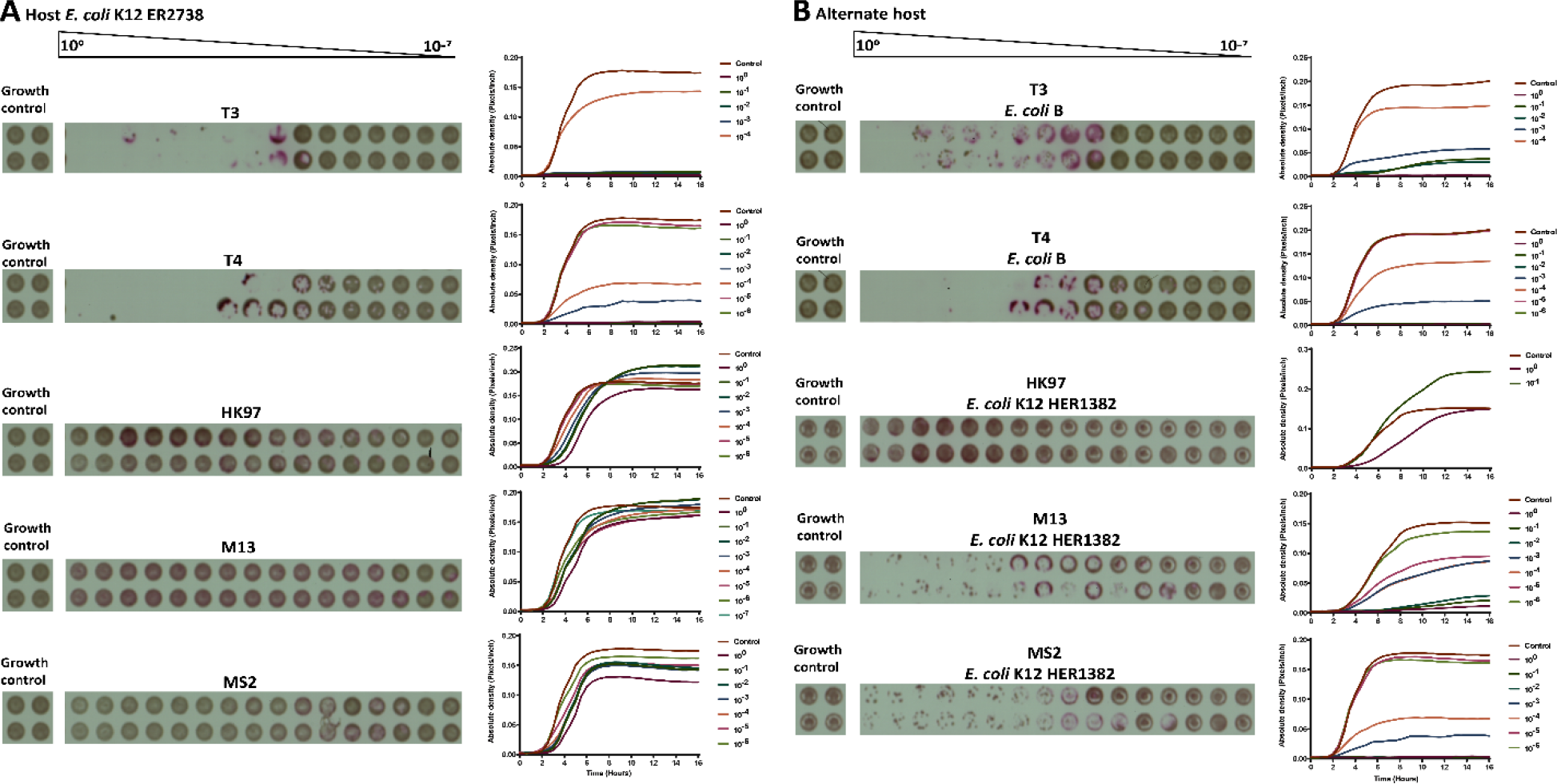
Biological replicate 2 of bacterial colonies with plaque formation of bacteriophages in micro-plaque assays. A) Left, image taken at endpoint for biological replicate 2. Each colony is represented by a tetrad of four technical replicates in a 10-fold dilution (10^0^-10^-7^). Growth controls for all phages was done using *E. coli* K12 HER2738 as host. Images were cropped from different plates according to the host, but not further manipulated. On the right side, plots represent the growth curves and their growth control, showing only curves in which plaques were counted, each curve represents the average of the four technical replicates per dilution. B) Same as A) but with different hosts as follows: T3 and T4 on *E. coli* B, with HK97 M13 and MS2 on *E. coli* K12 HER1382.

**Figure S3.**
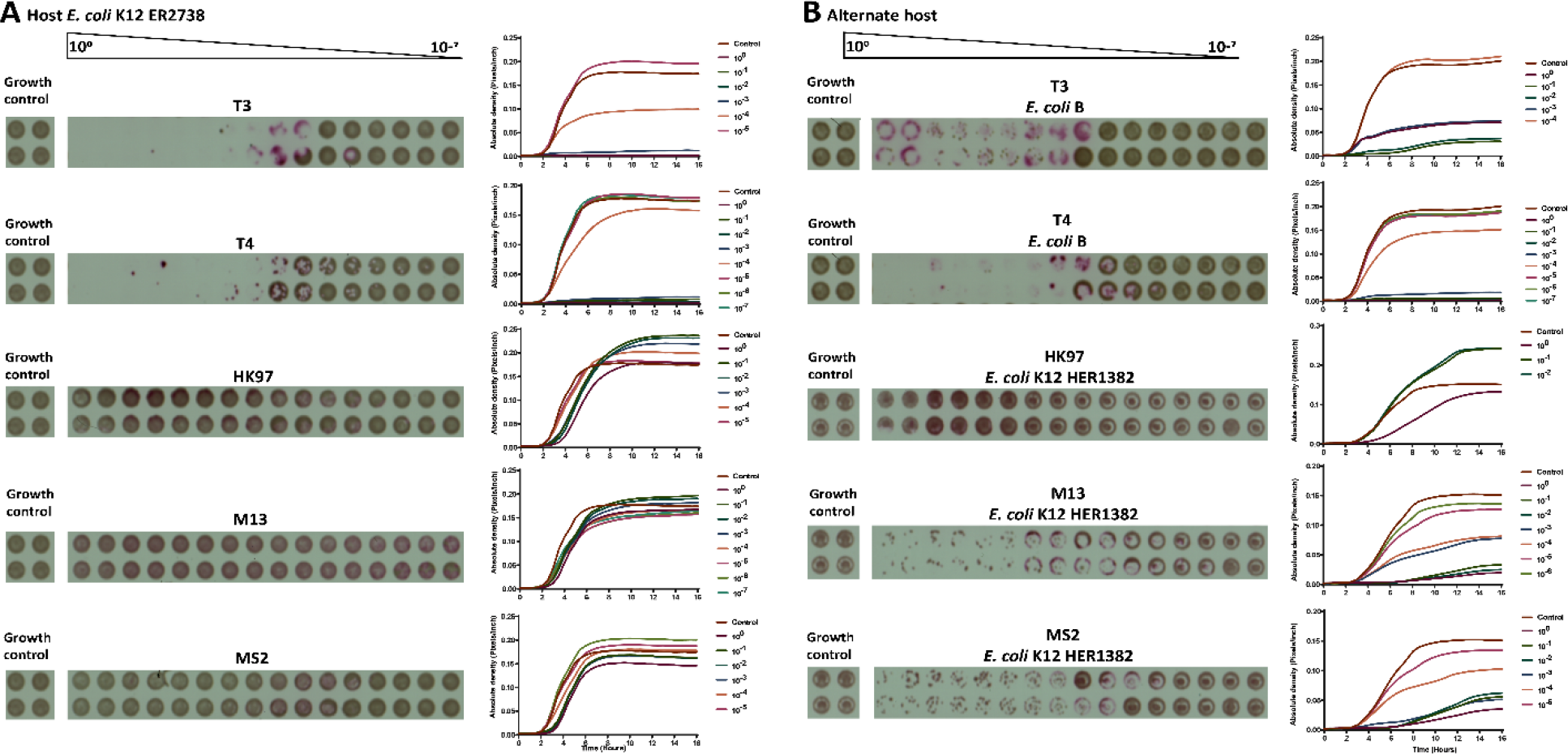
Biological replicate 3 of bacterial colonies with plaque formation of bacteriophages in micro-plaque assays. A) Left, image taken at endpoint for biological replicate 3. Each colony is represented by a tetrad of four technical replicates in a 10-fold dilution (10^0^-10^-7^). Growth control for all phages was done using *E. coli* K12 HER2738 as host. Images were cropped from different plates according to the host, but not further manipulated. On the right side, plots represent the growth curves and their growth control, showing only curves in which plaques were counted, each curve represents the average of the four technical replicates per dilution. B) Same as A) but different host as follow: T3 and T4 on *E. coli* B, with HK97 M13 and MS2 on *E. coli* K12 HER1382.

**Figure S4.**
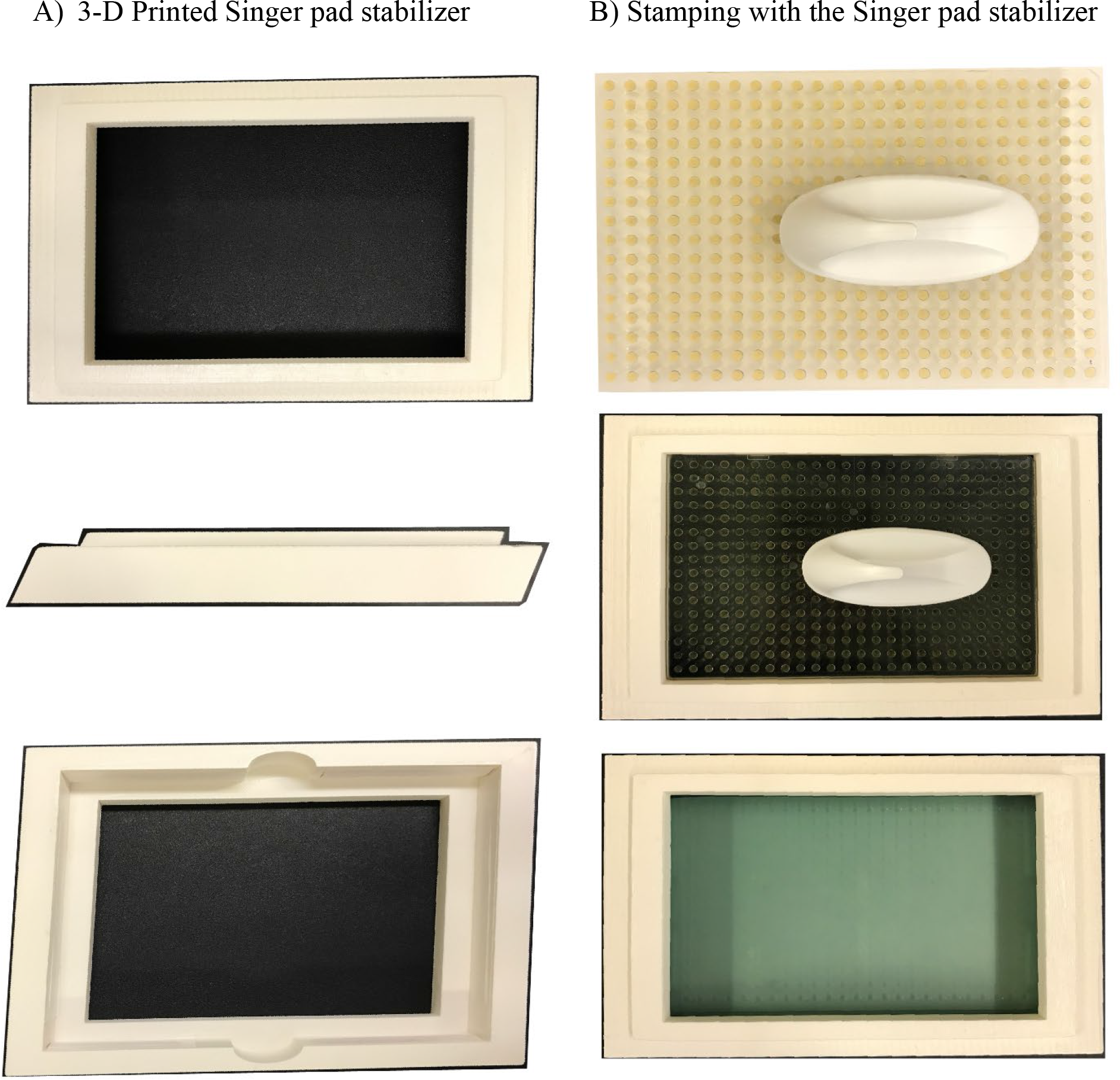
Visualization of the 3-D printed Singer pad stabilizer. A) From top to the bottom; images from each side of the 3-D printed Singer pad stabilizer. B) A Singer pad stabilizer with a pad placed inside to perform micro- plaque assays at 384-density.

**Figure S5.**
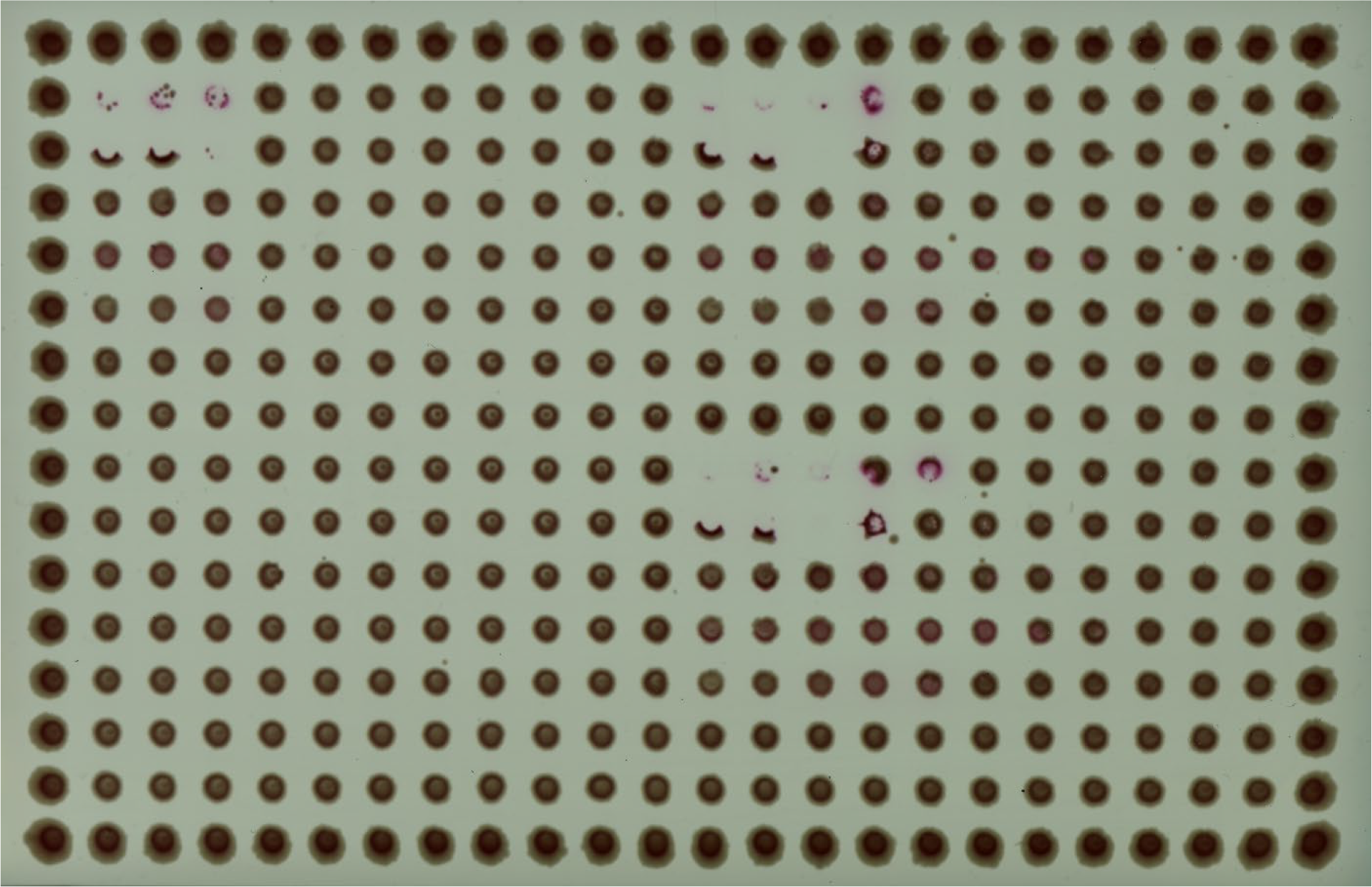
Manual micro-plaque assay at 384-density. Image taken at end point for three biological replicates of all phages on *E. coli* K12 HER2738. Each colony is represented by three technical replicates in a ten-fold dilution (10^0^-10^-7^). Image taken at end point and represents only one of the four technical replicates per dilution.

**Figure S6:**
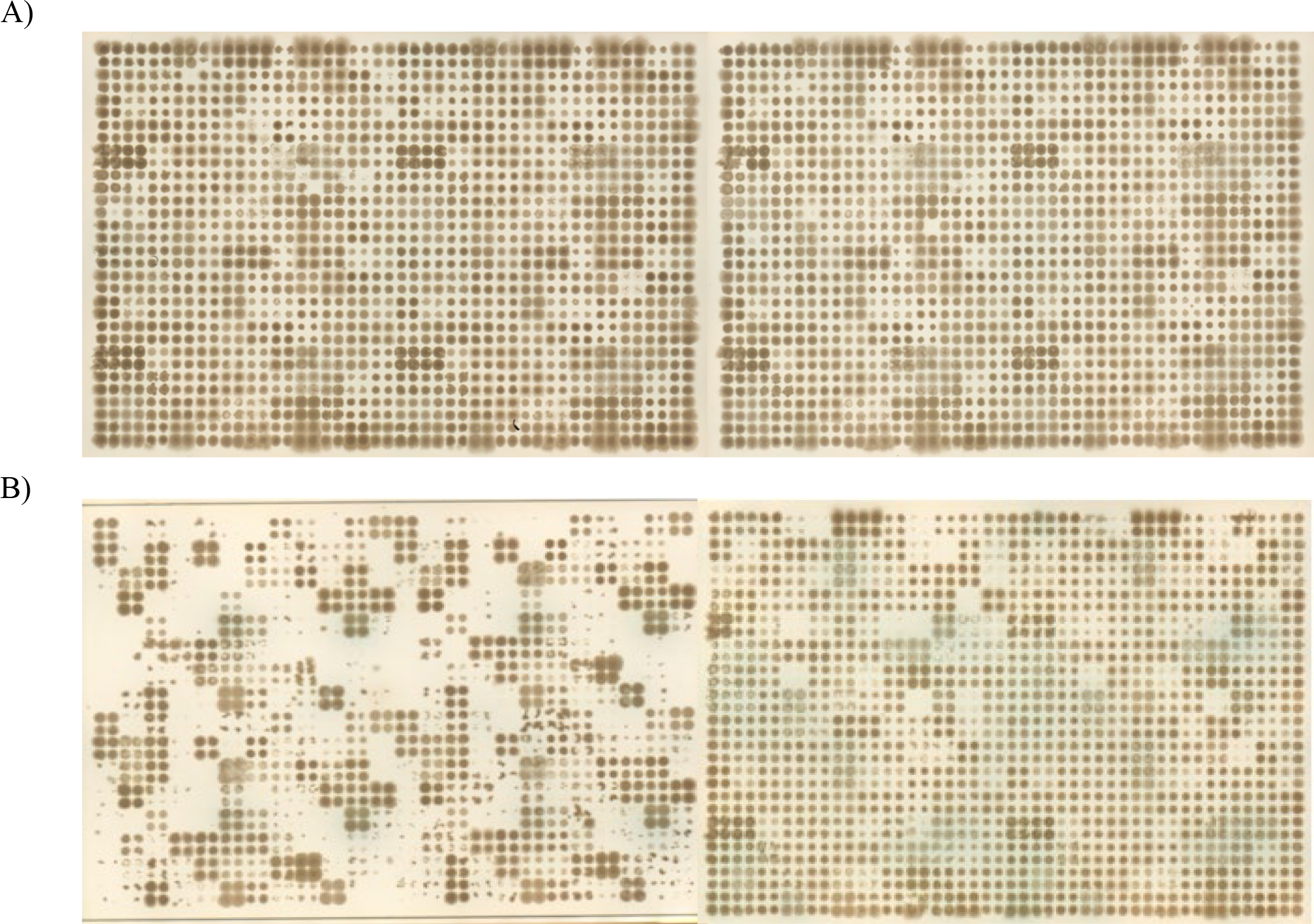
Reproducibility of the pipeline. A) Images illustrate a single isolated plaque generating similar phingerprints. B) Images illustrate two isolated plaques generating different phingerprints.

**Figure S7.**
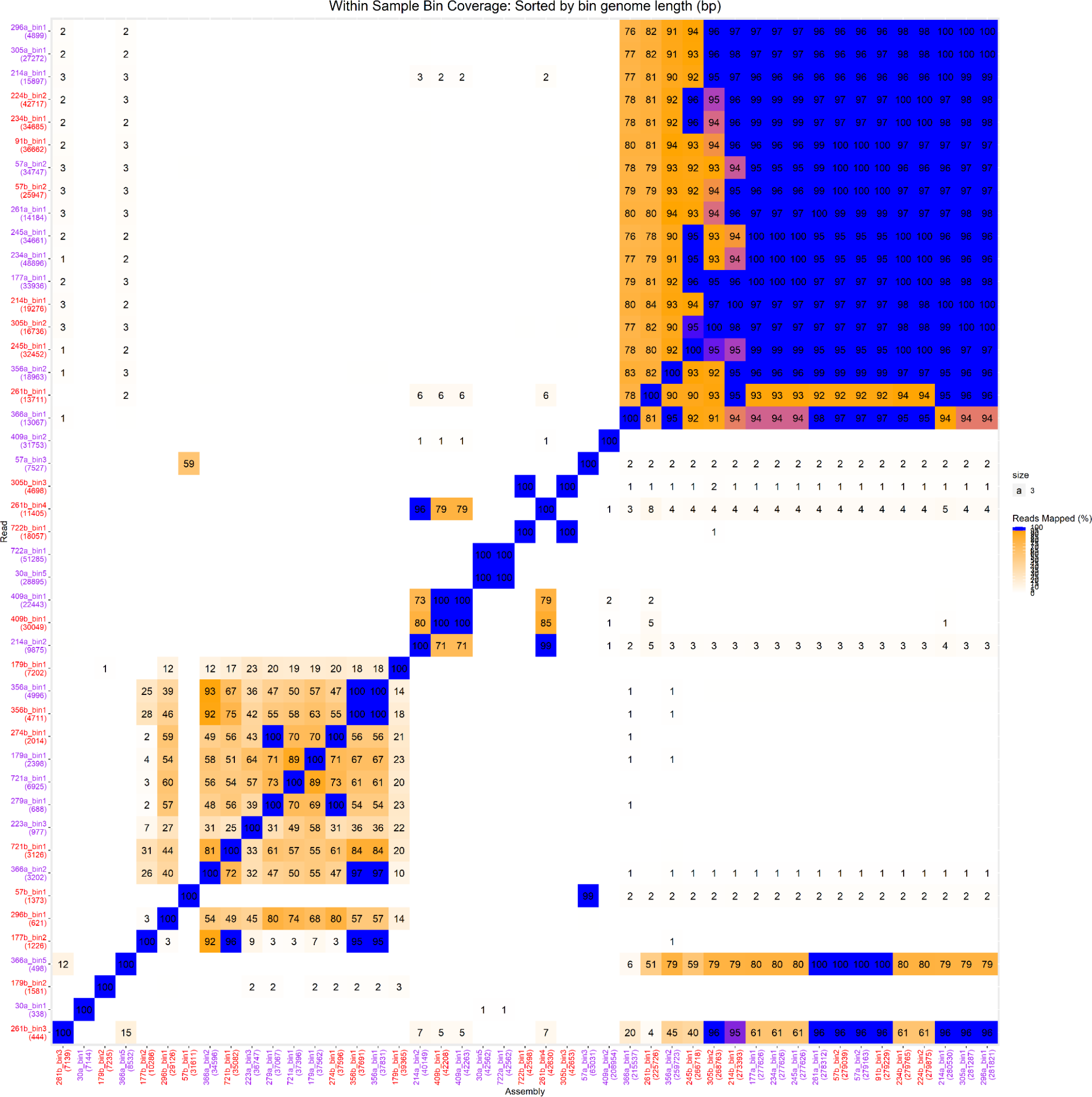
A summary of the comparison of all the genomic reads and assemblies between all samples of batch A and batch B. A total of 25 sample bins from batch A and 21 sample bins from batch B. Samples from batch A and batch B are distinguished by the purple and red colours respectively along each axis. The numeric value below each sample along the axis describes the total genomic length in base pairs (bp) ordered shortest (0,0 origin) to longest (0,1 or 1,0). The colour of each compared sample within the heatmap outlines the genomic similarity. The more similar a sample is to its counterpart the more blue the box. Any sample with a similarity less than 90% will appear as a yellow gradient colour. Yellow represents a low similarity and increases to a blue colour for highest similarity. The blue gradient begins at 90% similarity and continues to 100%. Each sample compared to itself is shown as a blue (100%) box forming the center diagonal line in the heatmap.

**Figure S8:**
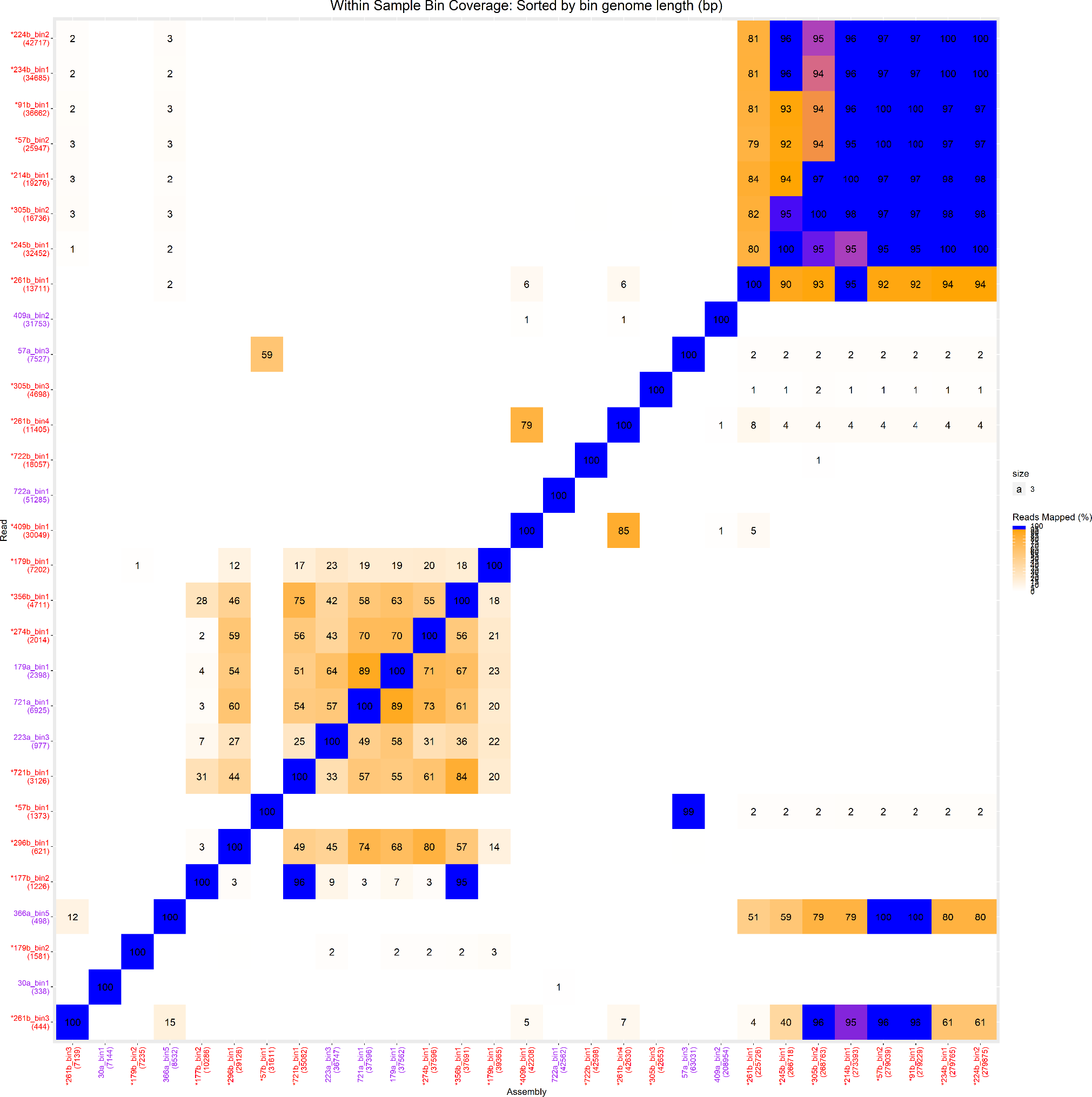
**A summary of the comparison of all the genomic reads and assemblies between only unique phages across batch A and batch B**. A subset of the total all against all comparison of batch A and batch B to contain only samples that were identified to be unique phage bins across the two batches. A total of 8 sample bins from batch A and 21 sample bins from batch B. Samples from batch A and batch B are distinguished by the purple and red colours respectively along each axis. The numeric value below each sample along the axis describes the total genomic length in base pairs (bp) ordered shortest (0,0 origin) to longest (0,1 or 1,0). The colour of each compared sample within the heatmap outlines the genomic similarity. The more similar a sample is to its counterpart the more blue the box. Any sample with a similarity less than 90% will appear as a yellow gradient colour. Yellow represents a low similarity and increases to a blue colour for highest similarity. The blue gradient begins at 90% similarity and continues to 100%. Each sample compared to itself is shown as a blue (100%) box forming the center diagonal line in the heatmap. The samples with an asterisk underwent additional downstream phylogenetic analysis.

**Figure S9:**
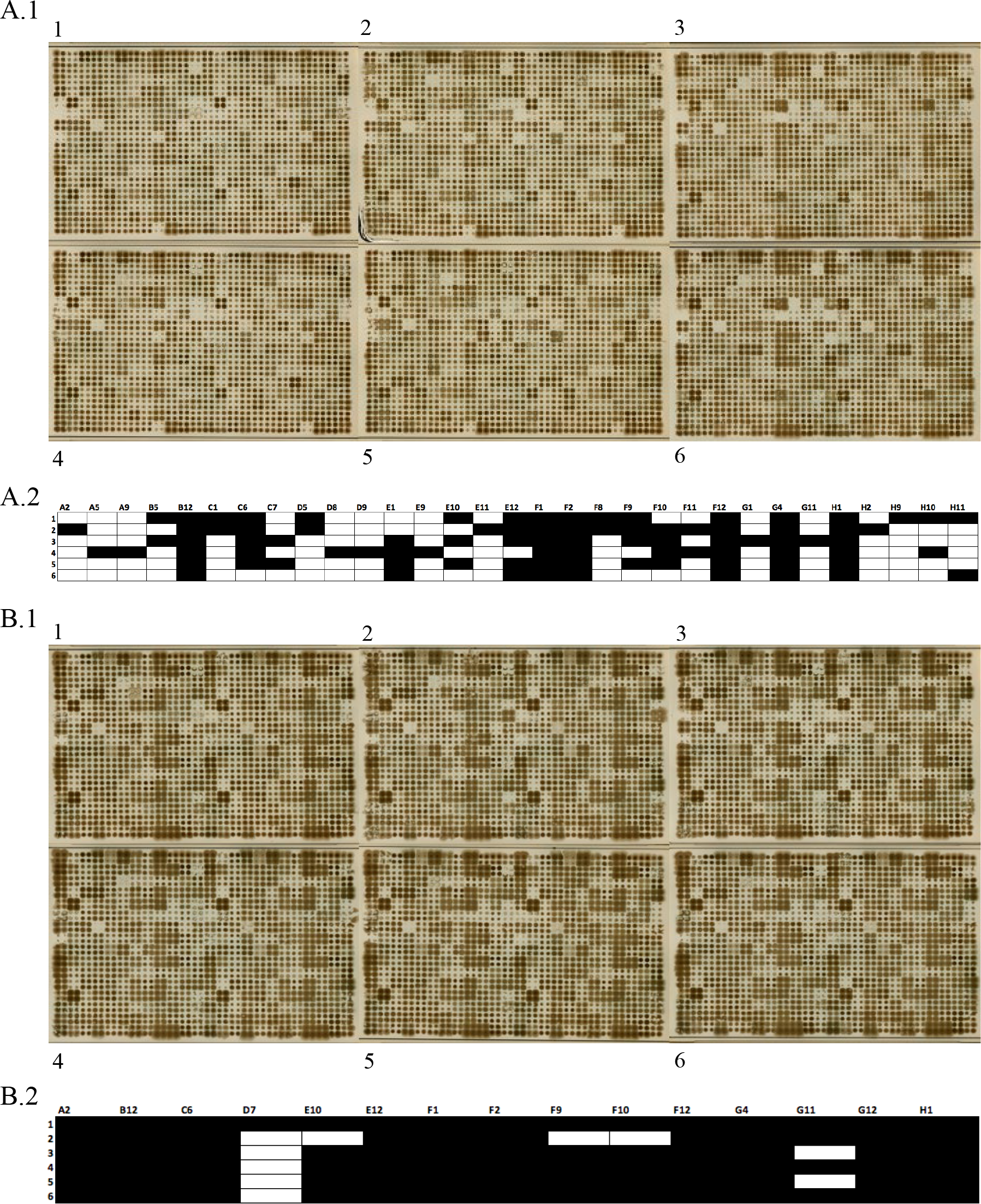
Amplification of phages on the same host display host range convergence. A.1) Phingerprint images of 6 phages amplified on bacterial host strains from which they were initially isolated during the phind pipeline. A.2) Corresponding host range of phages illustrated in A.1. Image portrays a binary presence (black)/absence (white) sequence indicating ability to lyse. Each column represents a different bacterial strain. B.1) Phingerprint images of the same 6 phages shown in A.1 amplified on the same bacterial host strain. B.2) Corresponding host range of phages illustrated in B.1. Image portrays a binary presence (black)/absence (white) sequence indicating ability to lyse. Each column represents a different bacterial strain.

**Table S1.** *E. coli* bacterial strains and phages for validation of the assays using the Singer Rotor HDA.

| Bacterial strain/phage family | Source |
| --- | --- |
| *E. coli* K12 ER2738 | 1 |
| *E. coli* K12 HER1382 | 2 (HER1382) |
| *E. coli* B | 2 (HER1024) |
| T3/Podoviridae | 2 (HER26) |
| T4/Myoviridae | 2 (HER27) |
| HK97/Shipoviridae | 2 (HER382) |
| M13/Inoviridae | 1 |
| MS2/Leviviridae | 3 |

1, NEB. 2, Félix D'Herelle Reference Centre for Bacterial Viruses. 3, DSMZ

**Table S2.** Efficiency of plaquing of micro-plaque assays versus standard plaque assays.

| Phage | Host | PFU/mL micro-plaque assay | EOP micro-plaque assay (*p*-value) | PFU/mL plaque assay | EOP plaque assay (*p*-value) | *p*-value across methods |
| --- | --- | --- | --- | --- | --- | --- |
| T3 | *E. coli* K12 ER2738 | 1.42x10<sup>8</sup>±79.6 | 0.8 (0.69) | 2.79x10<sup>8</sup>±121.4 | 1.4 (0.06) | 0.54 |
| T3 | *E. coli* B | 1.08x10<sup>8</sup>±70.5 |  | 4.02x10<sup>8</sup>±58.4 |  |  |
| T4 | *E. coli* K12 ER2738 | 5.83x10<sup>10</sup>±65.5 | 0.6 (0.62) | 2.25x10<sup>10</sup>±55.0 | 0.8 (0.61) | 0.20 |
| T4 | *E. coli* B | 3.75x10<sup>10</sup>±144.4 |  | 1.80x10<sup>10</sup>±37.1 |  |  |
| HK97 | *E. coli* K12 ER2738 | 2.50x10<sup>9</sup>±52.0 | 0.0001<br>**(0.03)** | 3.49x10<sup>8</sup>±13.8 | 0.002<br>**(0.0002)** | **0.046** |
| HK97 | *E. coli* K12 HER1382 | 2.50x10<sup>5</sup>±10.0 |  | 8.52x10<sup>5</sup>±10.7 |  |  |
| M13 | *E. coli* K12 ER2738 | 1.23x10<sup>11</sup>±108.8 | 0.1 (0.24) | 4.55x10<sup>10</sup>±79.6 | 1.5 (0.62) | 0.39 |
| M13 | *E. coli* K12 HER1382 | 1.50x10<sup>10</sup>±72.6 |  | 6.84x10<sup>10</sup>±95.8 |  |  |
| MS2 | *E. coli* K12 ER2738 | 9.42x10<sup>9</sup>±143.8 | 6.4 (0.33) | 5.11x10<sup>9</sup>±12.7 | 3.6<br>**(0.013)** | 0.61 |
| MS2 | *E. coli* K12 HER1382 | 6.00x10<sup>10</sup>±131.0 |  | 1.84x10<sup>10</sup>±29.3 |  |  |

Titres shows the average of three biological replicates and ± relative standard deviation in each method. EOP, efficiency of plaquing. EOP was calculated dividing the titres from alternate host by the titres obtained on *E. coli* K12 ER2738. Numbers in brackets indicate t-test *p*-values between titres from two groups. Numbers in bold indicates *p*-values <0.05.

**Table S3.**
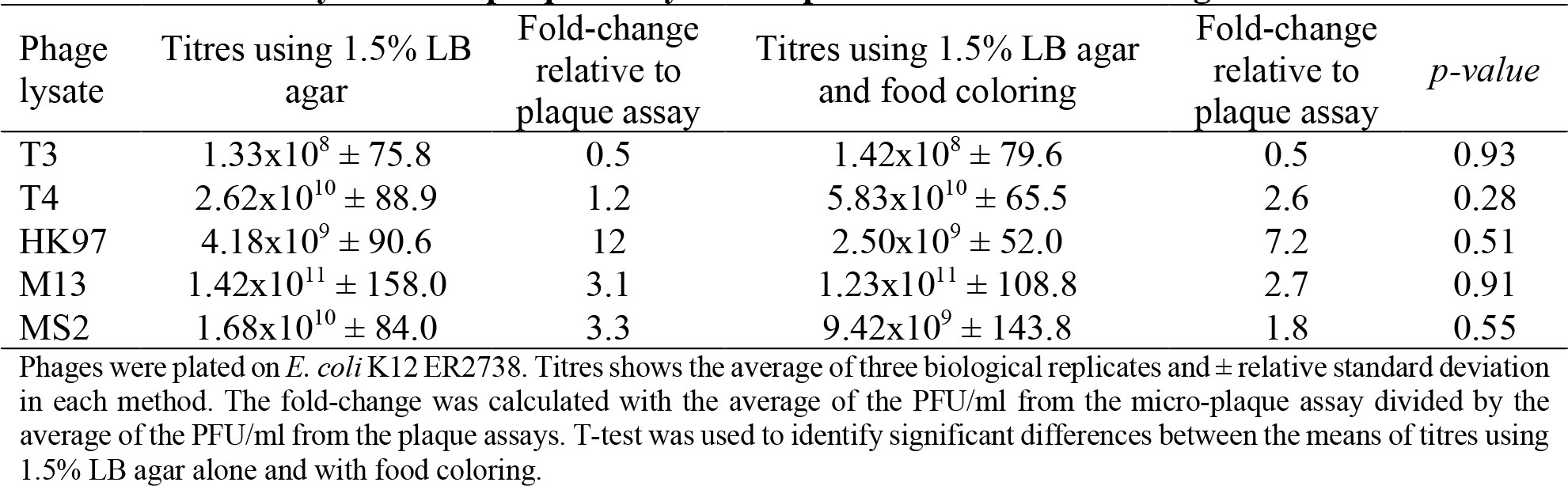
Sensitivity of micro-plaque assays in the presence of food colouring.

**Table S4.** Sensitivity of micro-plaque assays using 3D-printed Singer pad stabilizer at 384-density.

| Phage lysate | Titres using the Singer Rotor HDA | Fold-change relative to plaque assay | Titres using 3D-printed Singer pad stabilizer | Fold-change relative to plaque assay | *p-value* |
| --- | --- | --- | --- | --- | --- |
| T3 | $1.19 \times 10^8 \pm 102.7$ | 0.4 | $2.03 \times 10^8 \pm 128.3$ | 0.7 | 0.64 |
| T4 | $1.83 \times 10^{10} \pm 63.0$ | 0.8 | $5.01 \times 10^{10} \pm 99.6$ | 2.2 | 0.34 |
| HK97 | $2.58 \times 10^{09} \pm 92.0$ | 7.4 | $2.08 \times 10^9 \pm 124.8$ | 6.0 | 0.82 |
| M13 | $1.08 \times 10^{11} \pm 48.0$ | 2.4 | $4.25 \times 10^{10} \pm 120.1$ | 0.9 | 0.19 |
| MS2 | $6.25 \times 10^{09} \pm 155.9$ | 1.2 | $1.00 \times 10^9 \pm 131.5$ | 0.2 | 0.41 |

Phages were plated on *E. coli* K12 ER2738. Titres shows the average of three biological replicates and ± relative standard deviation in each method. The fold-change was calculated with the average of the PFU/ml from the micro-plaque assay divided by the average of the PFU/ml from the plaque assays. T-test was used to identify significant differences between the means of titres between two groups.

**Table S5.1.**
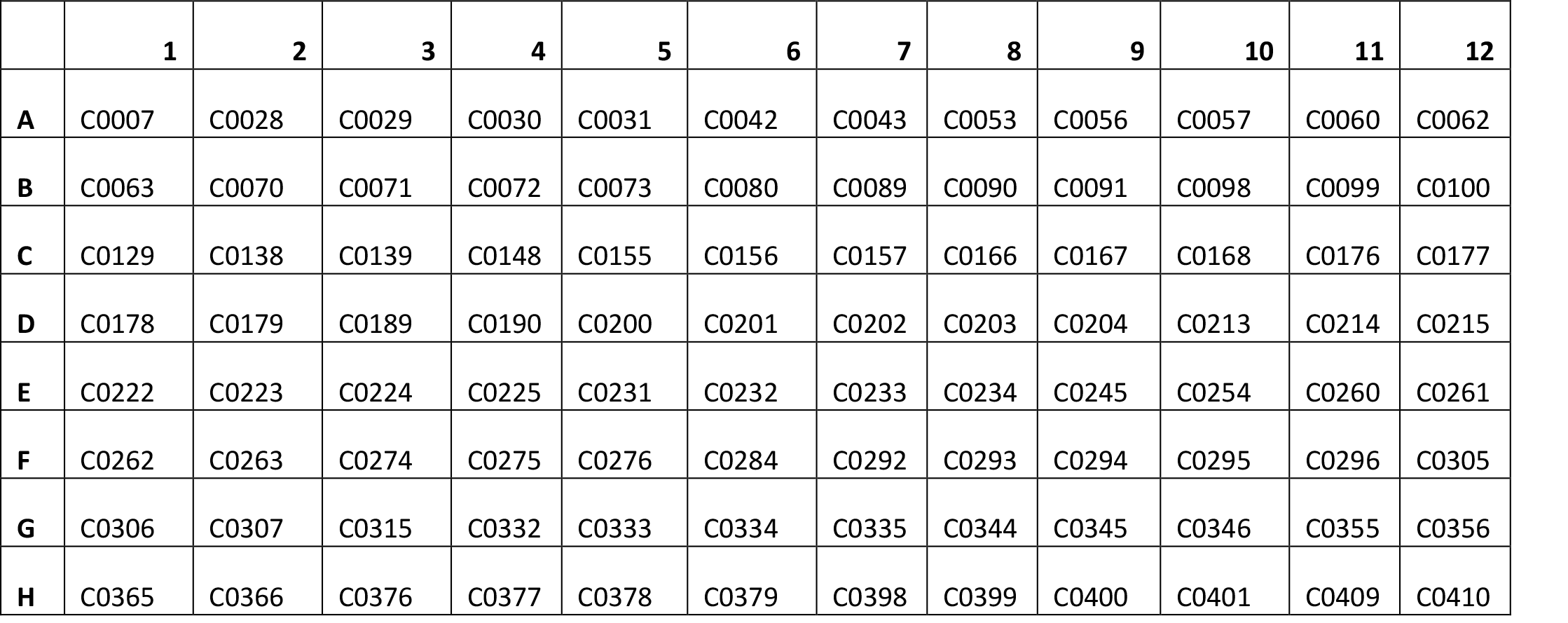
*P. aeruginosa* bacterial strains used for the phind assays.

**Table S5.2.**
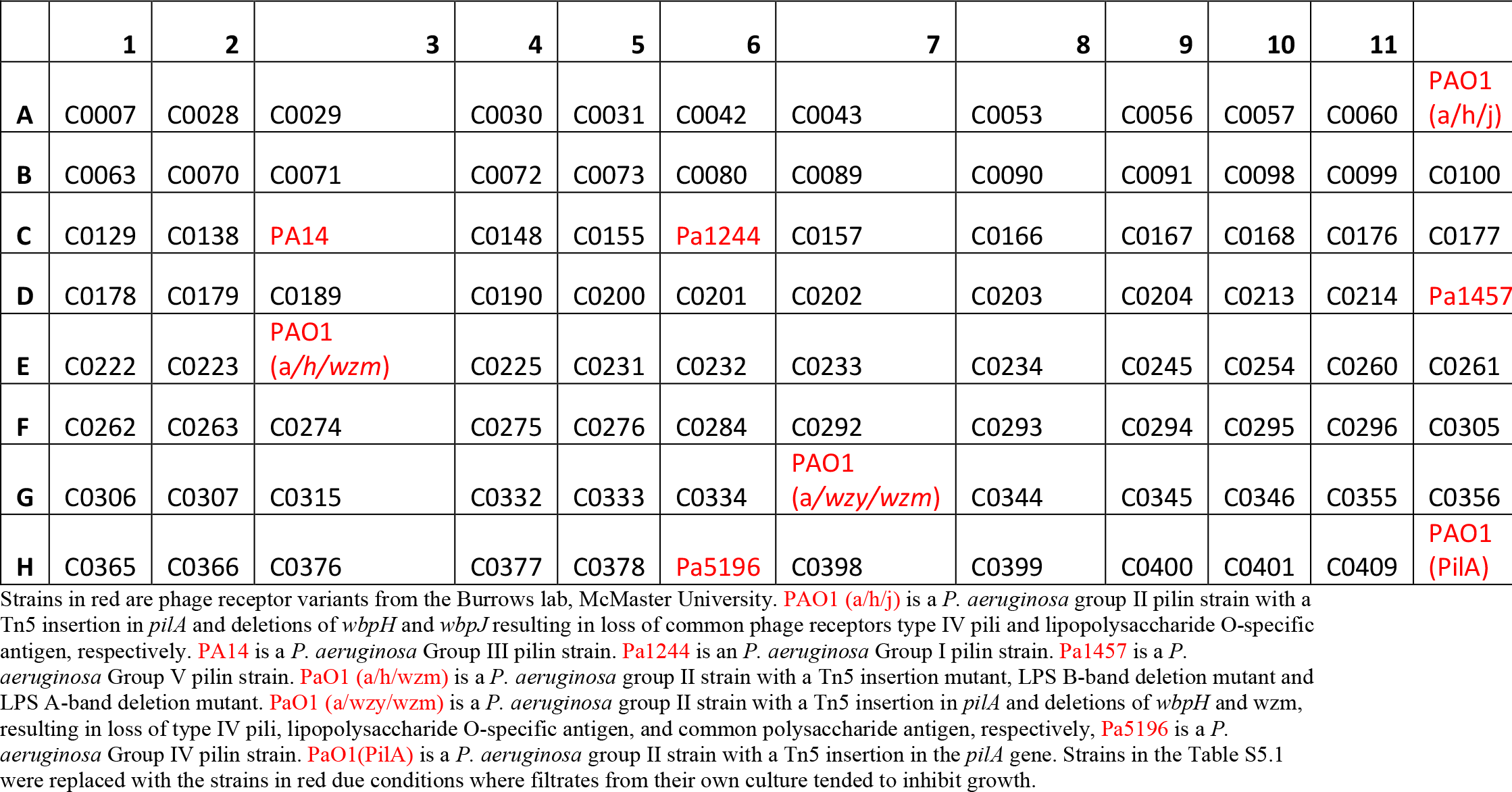
*P. aeruginosa* bacterial strains used for the phingerprint assays.

